# Network topology creates independent control of G_2_-M from G_1_-S checkpoints in the fission yeast cell cycle system

**DOI:** 10.1101/2025.03.09.642024

**Authors:** Yuhei Yamauchi, Hironori Sugiyama, Yuhei Goto, Kazuhiro Aoki, Atsushi Mochizuki

## Abstract

Physiological functions of cells arise from the dynamics of chemical reaction networks. The cell cycle of fission yeast is controlled by dynamical changes in two cyclin-dependent kinase (CDK)-cyclin complexes based on a complicated reaction network consisting of protein synthesis, complex formation, and degradation^1,2^. Each of the two checkpoints, G_1_-S and G_2_-M, is driven by an increase in the concentration of CDK-Cig2 and CDK-Cdc13, respectively. However, it is not understood how these complexes in the single connected network are controlled independently in a stage-specific manner. Here we theoretically predict that independent control of CDK-Cdc13 from CDK-Cig2 is achieved by the topology of the cell cycle network, and experimentally validate this prediction, while updating the network information by comparing predictions and experiments. We analyzed a known cell cycle network using a topology-based theory^3–6^ and revealed that the two CDK-cyclin complexes are included in different “regulatory modules”, suggesting that the concentration of each CDK-cyclin complex is controlled independently from the other. Experimental validation confirmed that the concentration of CDK-Cdc13 is controlled by the Cdc13 synthesis rate, independently from CDK-Cig2, as predicted. Conversely, the Cig2 synthesis rate affected not only CDK-Cig2 but also CDK-Cdc13. The fact, however, indicates the necessity of updating the network. We theoretically predicted the existence of an unknown necessary reaction, a Cdc13 degradation pathway, and experimentally confirmed it. The prediction and validation approach using the topology-based theory proposes a new systems biology, which progresses by comparing network structures with manipulation experiments and updating network information.

---

A large number of chemical reactions occurring within living cells are interconnected by sharing substrates and products, forming chemical reaction systems^7–10^. The rate of each reaction is controlled by enzyme activity, and by modulating the activity of each enzyme, organisms change the concentration of chemicals and thereby control physiological functions of cells^11–13^. The dynamics of chemicals based on chemical reaction systems are the origin of various physiological functions, such as metabolic homeostasis and cell cycle control, and each of these functions is achieved through the dynamical behaviors of multiple chemicals. A physiological function must be appropriately controlled in response to changes in the external signals or environment. Ideally, different functions should be independently controllable from each other. However, it has not even been understood whether different physiological functions arising from a single connected network system can be controlled independently.

## Structural sensitivity analysis (SSA) and a buffering structure

We here propose to apply a mathematically proven theoretical framework called structural sensitivity analysis (SSA)^3^ to address this question. SSA predicts the qualitative responses of chemical concentrations (and reaction fluxes) in steady state upon a change in a parameter such as enzyme activity or amount (or a conserved quantity). Importantly, SSA is based only on the structure of the reaction network for prediction and requires no information on explicit reaction rate functions (see Methods). This makes SSA a robust and general strategy for analyzing various biological networks, where the details of the reaction rate functions are largely unknown^14^. In particular, when SSA predictions do not align with experimental results, we can confidently conclude that the network information does not accurately represent reality.

Furthermore, SSA provides another powerful tool: the concept of a “buffering structure”^4,5^. A buffering structure defines a substructure of the network in which the effects of an arbitrary perturbation are confined in it at steady state. The buffering structure is determined by the local topology of the reaction network. In particular, if the number of elements in a substructure satisfies the following equation: – (number of chemicals) + (number of reactions) – (number of cycles) + (number of conserved quantities) = 0, it is called a buffering structure. It was mathematically proven that the response to a change in a parameter within a buffering structure is confined to inside it at steady state^4,5^ (Fig. 1a and also see Methods). In other words, the buffering structure provides a new concept that any substructure of a reaction network acquires physiological functions as a “regulatory module” simply by satisfying the above topological conditions^6^.

**Fig. 1.**
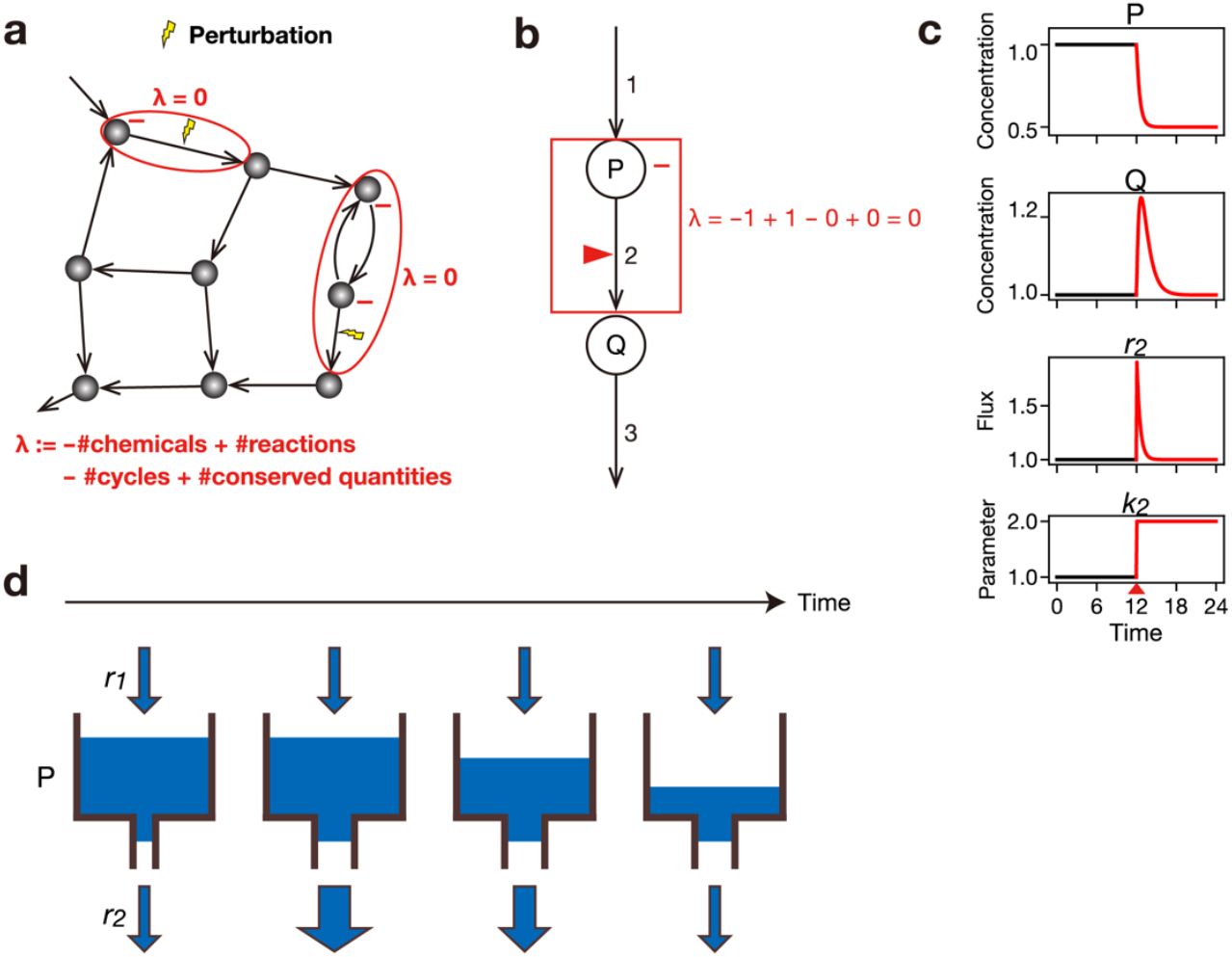
A buffering structure acts as a regulatory module in chemical reaction systems. **a**, Schematic of a buffering structure. A subnetwork that satisfies certain topological conditions—most importantly, the index λ ≔ − (number of chemicals) + (number of reactions) − (number of cycles) + (number of conserved quantities), is equal to 0—is called a buffering structure (See Methods for a detailed definition). The effects of a perturbation within a buffering structure are confined to inside the structure at steady state. **b**, Graphical representation of a chemical reaction system comprising of two chemicals (P,Q) and three reactions (1,2,3). Solid lines indicate chemical reactions. The red triangle indicates the reaction that is activated. The signs (increase/decrease) of responses are represented by +/− for chemicals, while chemicals without signs remain unchanged. The subnetwork enclosed by a red box is a buffering structure, with the index λ being equal to 0. **c**, A numerical simulation of the network in **b**. The concentrations of the chemicals (P, Q) and reaction rates of reaction 2 (*r*_2_) are plotted. Reaction rate functions and parameters are set as described in Methods. The dynamics are in the steady state at *t* = 12. At *t* = 12, *k*_2_, is increased by a factor of 2. **d**, Intuitive understanding of buffering structures using the water tank analogy (See the text for details).

For example, in the network shown in Fig. 1b, SSA shows that the steady-state concentration of chemical P decreases as a result of an increase in the enzyme activity catalyzing reaction 2 (Fig. 1b,c and Extended Data Fig. 1a,b). The steady-state concentration of chemical Q is not affected by this perturbation. These responses are understood by the buffering structure that encompasses reaction 2 and chemical P but not Q. Similarly, in the network shown in Extended Data Fig. 1c, SSA shows that an increase in the enzyme activity catalyzing reaction 3 induces a decrease in the steady-state concentration of W, and an increase in that of Z, while the steady-state concentrations of U and V do not change. These responses are understood by the buffering structure that encompasses reactions 3 and 4, and the chemicals W and Z (Extended Data Fig. 1c,d).

The function of a buffering structure can be understood intuitively as the behavior of a water tank with an inflow and an outflow. In Fig. 1d, increasing the enzyme activity of reaction 2 corresponds to enlarging the outlet. Then the outflow of water naturally increases, reducing the water level. The decrease in the water level induces a decrease in the water pressure at the bottom of the tank, which in turn decreases the outflow. This process stops when the inflow and outflow are in balance, realizing a new steady state, where the only net change is a decrease in the water level. In other words, the increase in the outlet size (enzyme activity) is compensated by the decrease in the water level (chemical concentration), and there is no change in the downward flow at steady state.

If a buffering structure exists in a network, it is theoretically predicted that chemicals inside it are independently controlled from chemicals outside. We hypothesize that the concept of buffering structures provides a theoretical basis for independent control of various functions in living organisms. In this paper, we demonstrate that this principle is actually at work in a real biological system.

## The cell cycle system

We choose the cell cycle system as a good example where multiple functions arise from a single connected reaction system. Progression through the cell cycle (G_1_/S/G_2_/M) is strictly controlled to ensure accurate genome replication and its subsequent distribution to the two daughter cells^15^. The G_1_-S and G_2_-M boundaries are called G_1_-S and G_2_-M checkpoints, respectively. Transitions through these checkpoints are driven by different cyclin-dependent kinase (CDK)-cyclin complexes. In the fission yeast *Schizosaccharomyces pombe*, increases in the concentration of CDK-Cig2 and CDK-Cdc13 trigger the transitions through the G_1_-S and G_2_-M checkpoints, respectively^16,17^ (Fig. 2a). These two CDK-cyclin complexes differ in their cyclin components (Cig2 and Cdc13) but share the same CDK (Cdc2). It has been thought that stage-specific control of the G_1_-S and G_2_-M checkpoints is achieved by the stage-specific increase in the expression of the two cyclins (Cig2 and Cdc13). That is, an increase in Cig2 expression would lead to an increase in CDK-Cig2 concentration, promoting the G_1_-S transition^2,17,18^, and an increase in Cdc13 expression would lead to an increase in CDK-Cdc13 concentration, promoting the G_2_-M transition^2,17,19^.

**Fig. 2.**
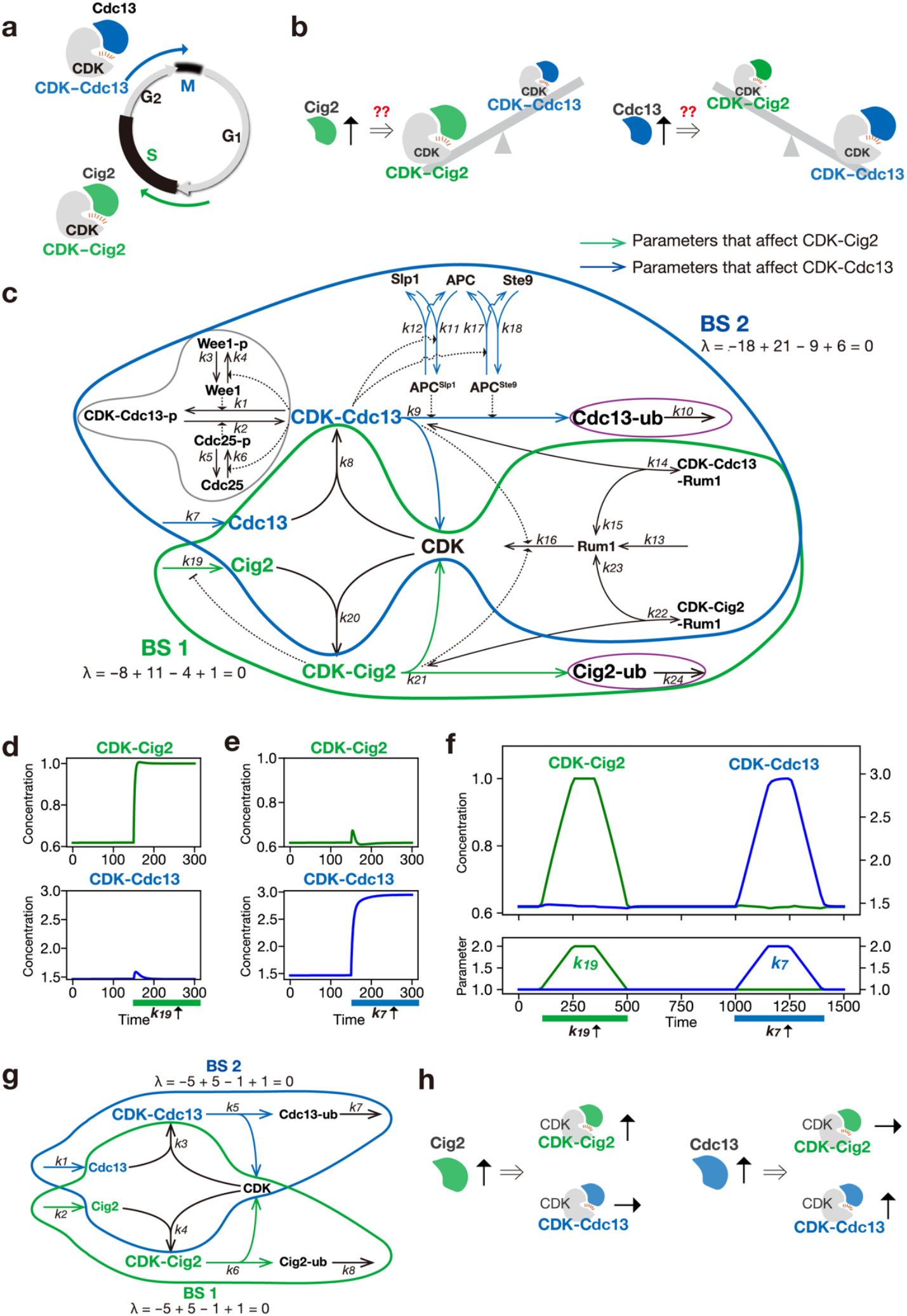
Analysis based on network structure demonstrates that CDK-Cig2 and CDK-Cdc13 are independently regulated. **a**, Schematic representation of the cell cycle system in fission yeast. **b**, As CDK-Cig2 and CDK-Cdc13 share the common CDK, it was intuitively hypothesized that these two complexes compete for CDK binding. **c**, A graphical representation of the cell cycle network. Each node indicates a chemical and each edge indicates a reaction (solid arc). Arrow-headed and line-headed dashed arc represents activating or inhibitory regulation, respectively. A reaction rate parameter (enzyme activity) for each reaction (*k*_*n*_) is depicted above or below the edge. Green arrows indicate reactions whose enzyme modulations affect the steady-state concentration of CDK-Cig2. Blue arrows indicate reactions whose enzyme modulations affect the steady-state concentration of CDK-Cdc13. The subnetwork enclosed by the green curve is a buffering structure containing CDK-Cig2 but not CDK-Cdc13 (BS1). The subnetwork enclosed by the blue curve is a buffering structure containing CDK-Cdc13 but not CDK-Cig2 (BS2). The intersection of BS1 and BS2 is also a buffering structure. The union of the intersection and the gray-enclosed region is another buffering structure, and the purple-enclosed regions are also buffering structures. **d**-**e**, Numerical calculation of the network dynamics are performed to illustrate the results in **c**. Reaction rate functions are assumed to be combinations of mass action and Michaelis–Menten kinetics. Initial values are set to a steady state of the system. At *t* = 150, the Cig2 synthesis rate (*k*_19_, **d**) or the Cdc13 synthesis rate (*k*_7_, **e**) is increased by a factor of 2. **f**, Numerical calculation of the network dynamics is conducted to recapitulate the cell cycle dynamics. We assume that the value of *k*_19_ or *k*_7_ change with time as shown below. **g**, A graphical representation of a simplified cell cycle network, consisting of cyclin synthesis reactions, CDK-cyclin complex formation and dissociation reactions, and cyclin degradation reactions. **h**, Summary of theoretical predictions based on the network structure.

As mentioned above, CDK-Cig2 and CDK-Cdc13 share the common CDK, and it is well known that the total amount of CDK remains almost constant throughout the cell cycle^20^. Based on this, it has been hypothesized that Cig2 and Cdc13 may compete for CDK binding^21^. Namely, when the expression level of Cig2 increases, not only CDK-Cig2 concentration will increase, but CDK-Cdc13 concentration might decrease. Conversely, if the expression level of Cdc13 increases, not only CDK-Cdc13 concentration will increase, but CDK-Cig2 concentration might decrease (Fig. 2b). However, the responses of the CDK-cyclin complexes to the changes in cyclin levels have not yet been confirmed quantitatively.

In this study, we used SSA and analyzed the cell cycle reaction system^1,2,22^ based on a currently known network structure. Contrary to the intuitive estimation based on the competition model, SSA predicted that the concentration of CDK-Cig2 and CDK-Cdc13 are independently controlled. We proved that this independent control is achieved by the structure of two characteristic buffering structures in the cell cycle network. That is, the buffering structure containing CDK-Cig2 (BS1) does not contain CDK-Cdc13. In other words, reaction parameters (enzyme amounts) in BS1 affect the concentration of CDK-Cig2, but never affect CDK-Cdc13. Similarly, the buffering structure that includes CDK-Cdc13 (BS2) does not include CDK-Cig2. In other words, reaction parameters (enzyme amounts) in BS2 affect the concentration of CDK-Cdc13, but never affect CDK-Cig2. If the network information is correct, such elegant, mutually exclusive control should actually be realized. In the following, we will elaborate on theoretical analysis of this system and experimental validations of the predictions. Different results were obtained for these two buffering structures through experimental validation, but both renew our understanding of the biological system.

## Structural sensitivity analysis (SSA) of the cell cycle network predicts independent control of CDK-Cig2 and CDK-Cdc13

Here, we study the network structure of the cell cycle system proposed in a previous study^2^ (Fig. 2c). This system consists of 20 chemicals and 24 reactions:

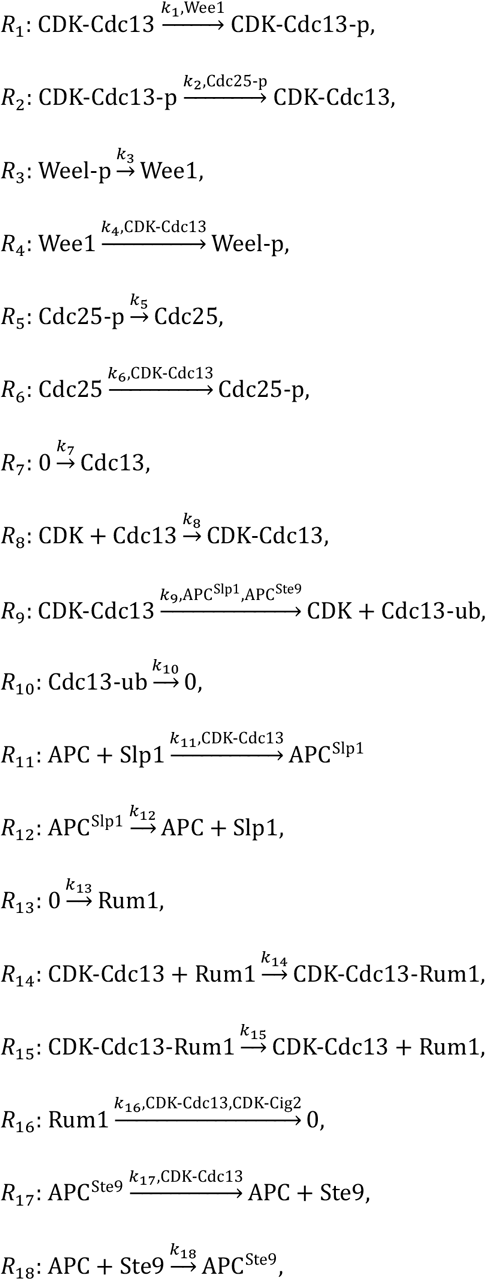

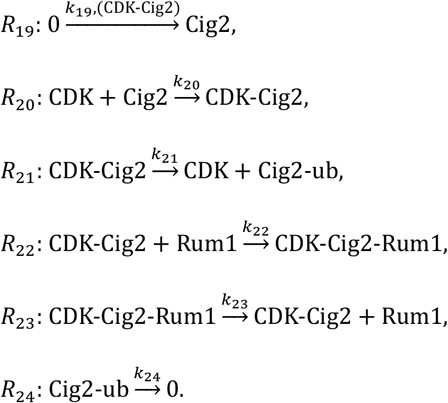

Above the arrow of each reaction *R*_*n*_, we display 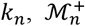 and 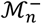.Here, *k*_*n*_ is the reaction rate parameter for *R*_*n*_, which can be the amount or the activity of the enzyme catalyzing the reaction. 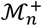 and 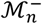 represent the sets of chemicals that are not substrates of the reaction *R*_*n*_ but regulate the reaction rate positively and negatively, respectively. 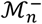 is indicated in parentheses and is present only for *R*_19_ (CDK-Cig2).

The activities of the enzymes catalyzing these reactions change as the cell cycle progresses. In particular, changes in the transcription and translation rates of cyclins (Cig2 and Cdc13) are considered the major driving force behind cell cycle progression^2,17–19^. In other words, the reaction parameters for Cig2 synthesis (*k*_19_) and Cdc13 synthesis (*k*_7_) are thought to change in a stage-dependent manner. While the yeast cell cycle takes around two hours, the reactions in the system progress in the timescale of seconds or minutes, and are much faster. Therefore, under given parameter values, it can be assumed that the system quickly reaches a steady state: temporal changes in the reaction parameters of cyclin synthesis (*k*_7_ or *k*_19_) are slow and thus are thought to cause shifts in this steady state.

Using SSA, we systematically identified the reaction parameters that affect the concentration of CDK-Cig2 or CDK-Cdc13 (Table 1 and Fig. 2c). We found that only a small number of reaction parameters can have an impact on these concentrations (Table 1 and Fig. 2c). For example, even if the synthesis rate of Rum1 (*k*_13_), which acts as a CDK inhibitor^23^, increases, it does not affect the concentration of any of the CDK-cyclin complexes. Similarly, changes in the reaction parameters associated with the phosphorylation/dephosphorylation reactions of Wee1 or Cdc25^24,25^ (*k*_3_, *k*_4_, *k*_5_, *k*_6_) do not affect any of the concentrations of CDK-cyclin complexes.

**Table 1.**
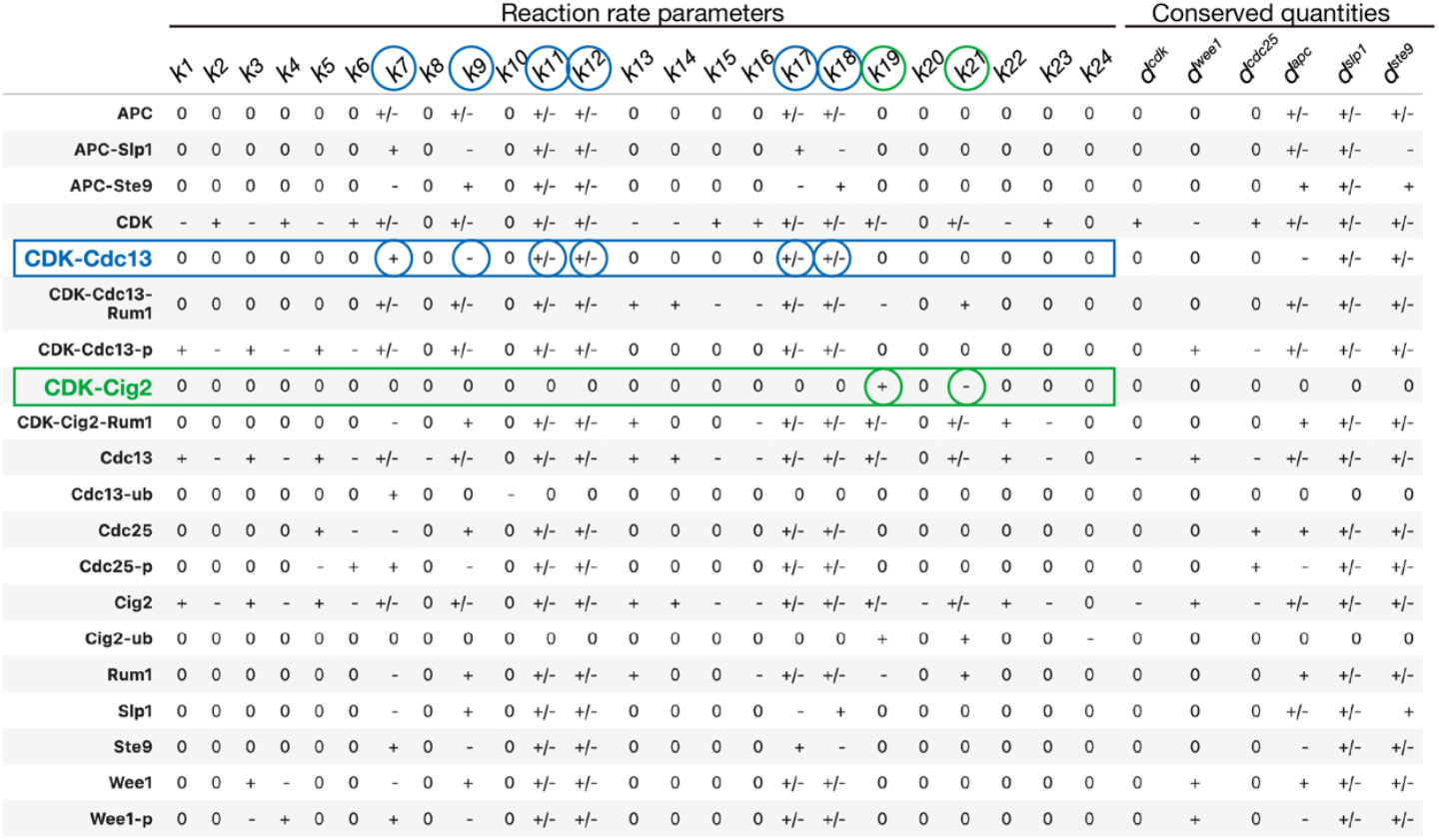
Structural sensitivity analysis (SSA) of the cell cycle network. A sensitivity matrix indicates qualitative changes in steady-state concentrations of each chemical (row) in response to the increase in each parameter (column), which is either reaction parameter or conserved quantity. The symbol “0” denotes no change, while the symbols “+” and “–” represent an increase and decrease, respectively. The symbol “+/–” indicates that the steady-state concentration changes, but the direction of the change cannot be determined solely from the network structure and instead depends on reaction rate functions or parameter values. Reaction parameters circled in green or blue represent parameters whose modulations alter the steady-state concentration of CDK-Cig2 or CDK-Cdc13, respectively.

Even more remarkably, there is no overlap between the reaction parameters affecting CDK-Cig2 and those affecting CDK-Cdc13 (Table 1 and Fig. 2c). In particular, increasing the synthesis rate of Cig2 (*k*_19_) raises the steady-state concentration of CDK-Cig2 but has no effect on that of CDK-Cdc13. Similarly, increasing the synthesis rate of Cdc13 (*k*_7_) raises the steady-state concentration of CDK-Cdc13 but does not affect that of CDK-Cig2. These results are different from the naïve prediction based on the competition model.

To illustrate the SSA prediction, we performed numerical simulations of the the cell cycle system dynamics by assigning mass action or Michaelis-Menten reaction rate functions to each reaction (Fig. 2d,e). During the simulation, we increased the Cig2 synthesis rate (*k*_19_) stepwise at a specific point. This perturbation caused CDK-Cig2 concentration to increase, and it converged over time to a new steady-state value higher than the original (Fig. 2d). In contrast, this perturbation caused a transient change in CDK-Cdc13 concentration, but it quickly returned to the original value, and the steady-state value did not change from before the perturbation (Fig. 2d). Similarly, when the Cdc13 synthesis rate (*k*_7_) was increased, CDK-Cdc13 concentration increased and converged to a new steady-state value higher than the original. (Fig. 2e). In contrast, CDK-Cig2 concentration temporarily increased, but quickly returned to its original value, and the steady-state value did not change from before the perturbation (Fig. 2e).

We also demonstrated how the concentrations of CDK-cyclin complexes change as the cell cycle progresses due to changes in reaction parameters (Fig. 2f). As the cell cycle progresses, the reaction parameter for Cig2 synthesis (*k*_19_) is thought to first increase and then decrease^2,17,18^; thus, we changed *k*_19_ accordingly. CDK-Cig2 concentration increased and decreased in response to the changes in the Cig2 synthesis rate, but CDK-Cdc13 concentration showed only a slight change. Similarly, the reaction parameter for Cdc13 synthesis (*k*_7_) is thought to change in the next stage of the cell cycle^2,17,19^; thus, we changed *k*_7_ in the later stage. During this phase, CDK-Cdc13 concentration increased and decreased in accordance with the changes in the Cdc13 synthesis rate, but CDK-Cig2 concentration only showed a slight change. In summary, if the known network information is correct, the concentrations of CDK-Cig2 and CDK-Cdc13 are controlled independently of each other (Fig. 2h).

## Independent control is achieved by distinctive buffering structures

We next aim to understand the mechanisms underlying independent control of CDK-Cig2 and CDK-Cdc13, specifically from the viewpoints of buffering structures (Fig. 2c). We identified two crucial buffering structures that play a pivotal role in the control of CDK-cyclin complexes. One is the buffering structure containing CDK-Cig2 but not CDK-Cdc13 (Fig. 2c and Extended Data Fig. 2a, green region, BS1), and the other is the buffering structure containing CDK-Cdc13 but not CDK-Cig2 (Fig. 2c and Extended Data Fig. 2a, blue region, BS2). BS1 indicates that any change in reaction parameters within BS1 does not affect the steady-state concentration of CDK-Cdc13, while influencing that of CDK-Cig2. For example, since the reaction parameter for Cig2 synthesis (*k*_19_) is contained within BS1, activating transcription and translation of Cig2 cannot affect the steady-state concentration of CDK-Cdc13, consistent with the SSA results. Similarly, BS2 indicates that any parameter change in BS2 does not affect the steady-state concentration of CDK-Cig2, while influencing CDK-Cdc13. For example, activating the Cdc13 synthesis rate (*k*_7_) cannot affect the steady-state concentration of CDK-Cig2.

All reactions in the system are contained within either of these two buffering structures. A mathematical proof guarantees that the intersection of these two buffering structures is also a buffering structure^26^. Since CDK-Cig2 and CDK-Cdc13 exist outside this intersection, the reactions within the intersection do not affect either of these two complexes. Within BS1 and BS2, we found other buffering structures that do not contain either CDK-Cig2 or CDK-Cdc13 (Fig. 2c, gray and purple regions), further narrowing down the set of reactions that influence either CDK-Cig2 or CDK-Cdc13. These findings indicate that independent control of the two complexes is achieved through two distinct buffering structures, one containing CDK-Cig2 and the other containing CDK-Cdc13.

## The effect of a change in a conserved quantity

In addition to changes in reaction rate parameters, SSA can also be used to analyze the response of a system caused by changes in conserved quantities within the system^5^. In this cell cycle network, the total concentration of CDK, i.e., the sum of free CDK and its binding complexes, is set to be constant over time (Extended Data Fig. 3a), consistent with a previous experimental report^20^. In reality, the concentration of this molecule may not be perfectly constant. To make sure that this uncertainty does not affect our conclusions, we performed the following analysis. Using SSA, we determined the qualitative changes in the concentration of each chemical when the total CDK amount changes. Among the six chemicals containing CDK, SSA predicted that only free CDK is affected by the increase in the total CDK amount, while the other five, including CDK-Cig2 and CDK-Cdc13, remain unaffected. The results of the numerical simulation also confirmed the SSA prediction (Extended Data Fig. 3b). This result is attributable to the existence of a buffering structure that includes CDK but not the other five CDK-containing complexes (Extended Data Fig. 3c). This buffering structure is a part of the intersection of BS1 and BS2. This result can be rephrased as follows: although we may intuitively expect that an increase in the total CDK amount would upregulate both CDK-Cig2 and CDK-Cdc13, this is not the case at all; rather, neither CDK-Cig2 nor CDK-Cdc13 concentrations change. In other words, the amount of CDK expression cannot be a driving factor for either the G_1_-S or G_2_-M transitions.

## The simplified network shows the same behavior as the original network

To clarify the structural conditions of the network that enable independent control of multiple checkpoints, we constructed a simplified cell cycle system consisting only of cyclin synthesis, CDK-cyclin complex formation, and dissociation, and performed SSA (Fig. 2g). Similar to the original cell cycle system, we found that the parameters affecting the concentration of CDK-Cig2 and those affecting the concentration of CDK-Cdc13 did not overlap, indicating that these complexes are controlled independently of each other. This is because the simplified system retains the same BS1 (containing CDK-Cig2 but not CDK-Cdc13) and BS2 (containing CDK-Cdc13 but not CDK-Cig2) as the original system (Fig. 2g).

These two buffering structures universally emerge when a reaction system including three chemicals (A,B,C) satisfies the following conditions: (i) the total amount of chemical A is conserved; (ii) chemical B is synthesized, binds to A, subsequently dissociates from A, and is then degraded; (iii) chemical C is similarly synthesized, binds to A, subsequently dissociates from A, and is then degraded.

## Experimental perturbations confirm that CDK-Cdc13 is controlled independently from CDK-Cig2

Do the behaviors predicted theoretically from the network structure actually occur in the fission yeast cell cycle system? We examined the effects of various perturbations on the concentrations of CDK-cyclin complexes using fluorescence cross-correlation spectroscopy (FCCS)^27–29^ (see Methods).

We first tested the existence of BS2, which includes CDK-Cdc13 but not CDK-Cig2 (Fig. 2c, blue). If this buffering structure exists, the perturbation to *k*_7_ (the Cdc13 synthesis rate) should affect the concentration of CDK-Cdc13 but not CDK-Cig2 (Fig. 2e,h and Fig. 3a). Additionally, since this buffering structure contains the conserved quantity CDK, perturbing the total CDK amount should not affect CDK-Cig2 (Extended Data Fig. 3b,c and Fig. 3d). FCCS analysis showed that overexpression of Cdc13 increased the concentration of CDK-Cdc13 while the concentration of CDK-Cig2 did not change (Fig. 3b,c). In addition, increasing the total CDK amount did not change the concentration of CDK-Cig2 (Fig. 3e). These results indicates that BS2 exists and that CDK-Cdc13 can be controlled independently from CDK-Cig2 in real fission yeast cells.

**Fig. 3.**
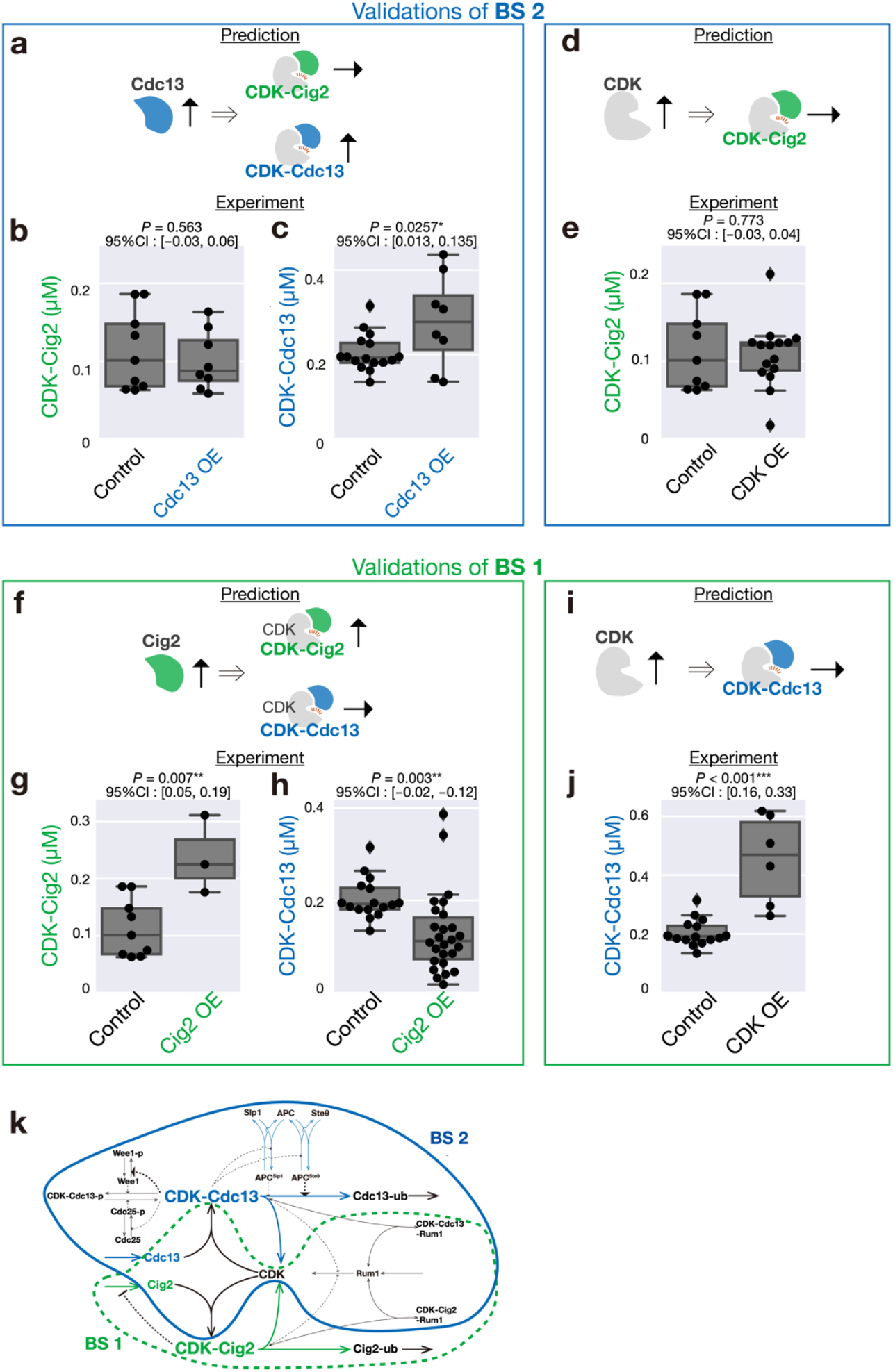
Experimental validation of theoretical predictions. **a**, Scheme of the theoretical predictions regarding the effects of Cdc13 overexpression on the concentrations of CDK-Cig2 and CDK-Cdc13. **b-c**, The concentrations of CDK-Cig2 (**b**) and CDK-Cdc13 (**c**) for control cells and Cdc13-overexpressing cells (Cdc13 OE). **d**, Scheme of a theoretical prediction regarding the effects of CDK overexpression on the concentration of CDK-Cig2. **e**, The concentration of CDK-Cig2 for control cells and CDK-overexpressing cells (CDK OE). **f**, Scheme of the theoretical predictions regarding the effects of Cig2 overexpression on the concentration of CDK-Cig2 and CDK-Cdc13. **g-h**, The concentrations of CDK-Cig2 (**g**) and CDK -Cdc13 (**h**) for control cells and Cig2-overexpressing cells (Cig2 OE). **i**, Scheme of a theoretical prediction regarding the effects of CDK overexpression on the concentration of CDK-Cdc13. **j**, The concentration of CDK-Cdc13 for control cells and CDK OE. **k**, Summary of experimental results. The box-and-whisker plots show the quartiles of the data, with whiskers extending to the minimum and maximum values, except for outliers, which are defined as values exceeding 1.5 times the interquartile range. \**P* < .05, \*\**P* < .01, \*\*\**P* < .001 and 95% confidence intervals (CI) of the differences (OE - Control) by Student’s *t* test.

## CDK-Cig2 is not independently controlled from CDK-Cdc13

We next tested the existence of BS1, which includes CDK-Cig2 but not CDK-Cdc13 (Fig. 2c, green). If this buffering structure exists, the perturbation to *k*_19_ (the Cig2 synthesis rate) should affect the concentration of CDK-Cig2 but not CDK-Cdc13 (Fig. 2d,h and Fig. 3f). Additionally, since this buffering structure contains the conserved quantity CDK, the perturbation to the total CDK amount should not affect CDK-Cdc13 (Extended Data Fig. 3b,c and Fig. 3i). FCCS analysis showed that overexpression of Cig2 upregulated CDK-Cig2 (Fig. 3g). However, it also led to a decrease in the concentration of CDK-Cdc13 (Fig. 3h). FCCS analysis also showed that increasing the CDK amount increased the concentration of CDK-Cdc13 (Fig. 3j). These findings show that BS1 does not exist in the fission yeast cell cycle system (Fig. 3k).

## Finding a new pathway by comparing predictions of the topology-based theory with experiments

Our theoretical predictions were derived from the network structure alone, without depending on any choices about reaction rate functions or parameters. If the network information is correct, the predicted behavior should always be observed. Therefore, the fact that some of the theoretical predictions are inconsistent with the experimental results means that the network information we used differs from the actual network. All the reactions included in the network are supported by sufficient experimental evidence^1,2,23–25,30^, so we are certain that there exist unidentified reactions that are not included in the current network information.

To identify the missing reactions, we conducted a theoretical search for the missing network information. The unidentified reaction must satisfy the following condition: by adding the reaction, BS1 is destroyed while BS2 is maintained. In general, for a buffering structure to be destroyed, a reaction (or regulatory) arrow must be added from the chemical inside the buffering structure to the outside^31^. In contrast, adding an arrow that goes into a chemical within a buffering structure never breaks the buffering structure unless the arrow breaks the conserved quantities. Therefore, here we only need to examine reaction arrows emanating from the chemicals within BS1 to the outside.

We introduced outgoing reactions for each chemical in the network and conducted SSA. Among the 13 possible network modifications, only one modification breaks BS1 while maintaining BS2. This modification is the addition of the degradation pathway for the Cdc13 monomer (Fig. 4a and also see Methods). We refer to the network including this modification as the “modified network” (Fig. 4b). In the original network, Cdc13 is assumed to be ubiquitinated only when it binds to CDK in an anaphase-promoting complex (APC/C)-dependent manner. In the modified network, Cdc13 that does not form a complex with CDK should be degraded. This breaks BS1 but not BS2, fully explaining the observed behavior (Extended Data Fig. 4). The same conclusion applies to the simplified cell cycle network: only the addition of a degradation reaction for the Cdc13 monomer breaks BS1 while maintaining BS2 (Fig. 4c and Extended Data Fig. 5).

**Fig. 4.**
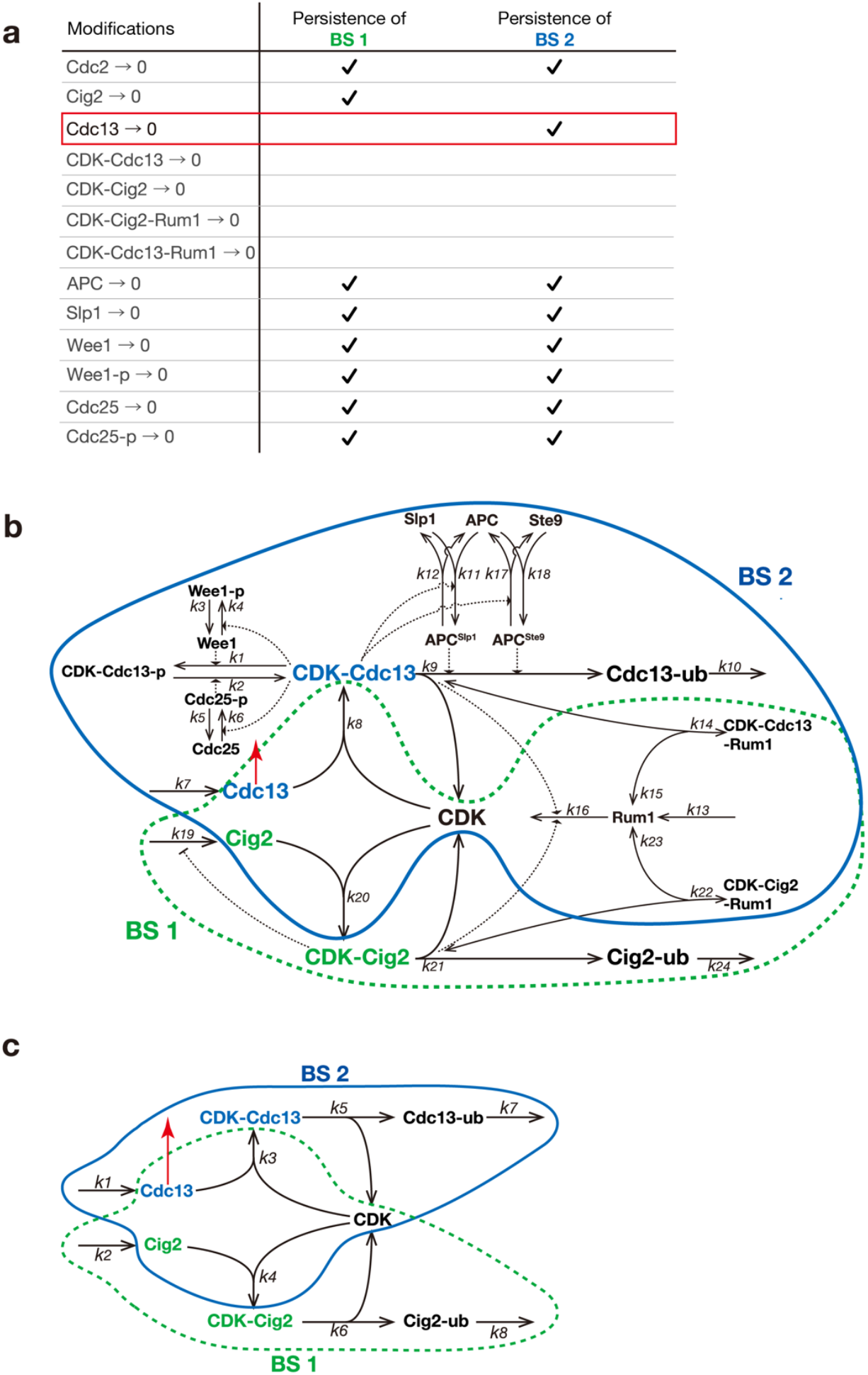
Theoretically identifying missing reactions in the cell cycle network. **a**, We considered network modifications where a degradation reaction is added to each of the chemicals that originally do not have a degradation reaction. Each modification was then evaluated for whether it satisfied the condition that BS1 is destroyed and BS2 is maintained. We find that only the addition of degradation of the free Cdc13, which is not bound to CDK, (Cdc13 →0) satisfies the condition, thereby explaining the experimental results. **b**-**c**, Graphical representations of the modified cell cycle network (**b**), and modified simplified network (**c**). A red reaction, representing the degradation of Cdc13 that does not bind to CDK, is added. This modification causes BS1 to disappear, while BS2 is maintained.

## Experimental validation of the predicted pathway

We finally examined whether this predicted pathway (degradation of the Cdc13 monomer) exists in actual fission yeast cells. Given that APC/C primarily degrades Cdc13 bound to CDK, as shown in Fig. 2c, the degradation of the Cdc13 monomer may not depend on APC/C. We thus hypothesized that an APC/C-independent Cdc13 degradation pathway exists. To test this hypothesis, we conducted a cycloheximide (CHX) chase analysis^19,32^ for *cdc13-mNeonGreen* cells, and cells expressing mNeonGreen not fused to Cdc13 (negative control). CHX is a translation inhibitor that blocks new protein synthesis, allowing us to monitor the degradation of proteins over time ^19,32^. After adding CHX, we measured the fluorescence intensity of mNeonGreen at various time points for each cell. Given that APC/C becomes active just before anaphase in M phase^30^, we used image analysis to distinguish cells before nuclear division (G_2_ phase) from those after nuclear division (M phase). (Fig. 5a). The degradation rate of Cdc13 in G_2_ phase was lower than that in M phase. However, the degradation rate of Cdc13 in G_2_ phase was still higher than that of the negative control (Fig. 5b,d,e), suggesting that Cdc13 is actively degraded in G_2_ phase, where APC/C is inactive. Consistent with this observation, pharmacological inhibition of APC/C did not decrease the degradation rate of Cdc13 in G_2_ phase (Fig. 5c,d,e). These results confirm the existence of an APC/C-independent Cdc13 degradation pathway, which was predicted by comparing the SSA expectations with experimental results.

**Fig. 5.**
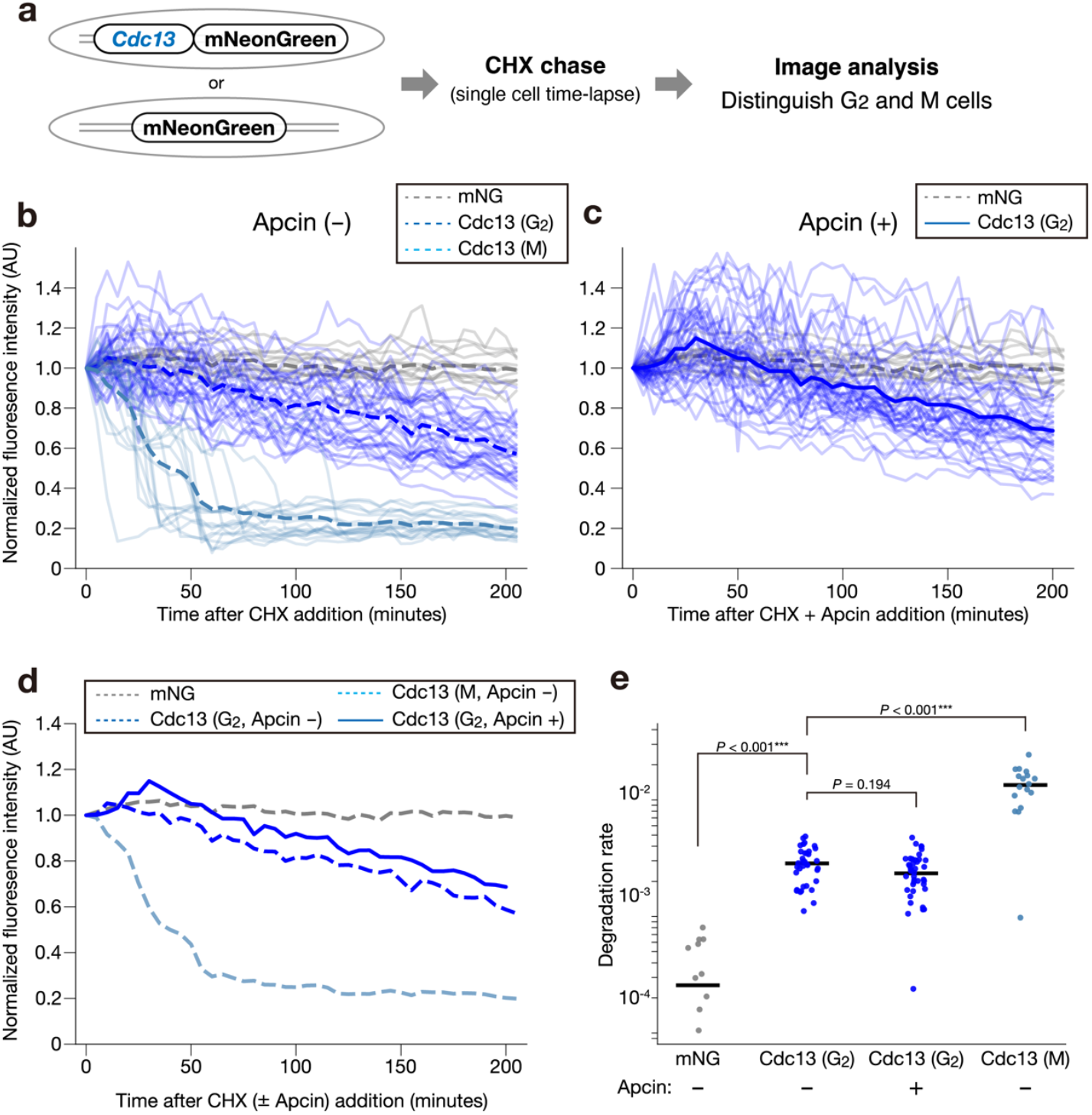
Cdc13 is degraded in non-M phase in an APC-independent manner. **a**, Schematic representation of the experimental strategy to measure the degradation rate of Cdc13. *Cdc13-mNeonGreen (mNG)* cells were used, and mNG not fused to Cdc13 was used as a control. Using image analysis, we distinguished cells that did not go through nuclear division (G_2_ cells) and cells that went through nuclear division (M cells). **b**, Time course for the relative fluorescence levels of control (gray) and Cdc13-mNG (blue) for G2 cells under the treatment of CHX. Cdc13-mNG for M cells (light blue) is also shown. CHX is added at time 0. Time-lapse images were acquired using confocal microscopy. N = 35 cells for G2 cells, N = 17 for M cells, and N = 13 for a control. **c**, Time course for the relative fluorescence levels of control (gray) and Cdc13-mNeonGreen (blue) under the treatment of CHX with the APC inhibitor Apcin. CHX and Apcin are added at time 0. N = 42 cells. **d**, Average time course for the mNG and Cdc13 levels in (**b**) and (**c**) are overlaid. **e**, The degradation rates of mNG or Cdc13 of each cell estimated by fitting the data in (**b**) and (**c**) to an exponential decay model. Horizontal bars are means. \*\*\**P* < .0001 by Student’s *t* test adjusted by Bonferroni method for multiple comparisons.

## Discussion

To clarify the regulatory mechanisms of the two checkpoints in the cell cycle, we analyzed the cell cycle network based solely on its structure using the topology-based theory SSA. Our analysis predicted that if the network information is correct, the concentrations of CDK-Cig2 and that of CDK-Cdc13 are controlled independently of each other. This independent control is achieved by the structure of the two characteristic buffering structures in the cell cycle network. That is, the buffering structure containing CDK-Cig2 but not CDK-Cdc13 (BS1), and the buffering structure containing CDK-Cdc13 but not CDK-Cig2 (BS2).

We validated this prediction by generating fission yeast mutants that overexpress Cig2 or Cdc13 and then measuring CDK-Cig2 and CDK-Cdc13 concentrations using FCCS. As a result, we found that overexpression of Cdc13 increased the concentration of CDK-Cdc13, but did not change that of CDK-Cig2, confirming the existence of BS2. Conversely, a change in Cig2 expression not only affected CDK-Cig2, but also caused a change in the concentration of CDK-Cdc13, indicating that BS1 does not exist.

Since the SSA predictions are derived from network information alone, the discrepancy between the theoretical predictions and actual behavior is justified only by two scenarios: the known network information is incorrect, or the validation experiment is wrong. We theoretically searched for all possible network modifications satisfying the condition that BS2 persists and BS1 vanishes. Only a single modification satisfies this criteria: the existence of degradation of Cdc13 monomer, which we confirmed experimentally.

In the following, we develop our discussion in three directions: 1) The elucidation of the cell cycle system, 2) topology-based theory, and 3) the combination between topology-based predictions and manipulation validation.

1) First, we clarified that the transition through the G_2_-M checkpoint is controllable independently from the G_1_-S checkpoint. This is an achievement of successful collaboration between theory and experiment: predicted by theoretical analysis of network structure and verified by precise quantitative measurements. As the two CDK-cyclin complexes governing the two checkpoints share the same CDK, the competition hypothesis has been proposed, which states that the two cyclins may always compete with each other for CDK binding. Our results refute this hypothesis, strongly suggesting a new scheme from both theoretical and experimental viewpoints. The fact that the G_2_-M checkpoint is controllable independently from the G_1_-S checkpoint might be important for cell function. The G_1_-S checkpoint and the G_2_-M checkpoint have different roles, determining the timing of the initiation of genome replication and that of mitosis, respectively. The fact that the control of the initiation of mitosis is independent from that of the initiation of replication is likely to be adaptive for proper cell function. In particular, even if the amount of Cdc13 inappropriately increases due to some expression fluctuation during the G_1_-S transition, it would never influence CDK-Cig2 and therefore normal G_1_-S transition would be realized. On the other hand, an irregular increase in Cig2 concentration not only affects CDK-Cig2 but also CDK-Cdc13. How should we understand this? In fission yeast, while the expression level of Cdc13 rapidly increases at a specific stage, Cig2 expression does not exhibit remarkable stage-specific changes^20^. Therefore, even if an irregular fluctuation occurs in Cig2 expression, the magnitude of the expression change would be small and its effect would be negligible. For these reasons, the selective pressure to ensure that Cig2 expression changes do not affect CDK-Cdc13 might not have been very strong. Applying SSA to the cell cycle systems of other species, including humans, is a potential direction for future research. Unlike fission yeast, mammalian cells have multiple CDKs, including CDK1, CDK2, CDK4, and CDK6. Among these, only CDK1—the counterpart to fission yeast’s Cdc2—is essential, since the cell cycle progresses almost normally in most cell types even in the absence of other CDKs^33–37^. CDK1 binds to all cyclins when other CDKs are absent^35^. This suggests that fundamental regulatory mechanisms governing the cell cycle are conserved between fission yeast and mammals. It is thus highly probable that buffering structures also play a crucial role in the mammalian cell cycle, just as they do in fission yeast.
2) Second, we have clearly shown that the mechanism of independent control arises solely from the structural properties of the reaction network. Many theoretical approaches have attempted to understand complex biological systems in terms of network structures^38,39^, but many of them require assumptions about specific functional forms, and cannot be said to be purely topology-based. We have developed several purely topology-based theories^3–6,31,40–46^, and SSA is one of them. The concept of a buffering structure mathematically predicts that network topology creates independent control of multiple functions emerging from a single connected network. This study clearly demonstrates, for the first time, the physiological link between the mathematical concept of a buffering structure and molecular basis that enables independent control of interconnected systems in living cells. To clarify the structural conditions of the network that enable independent control of multiple checkpoints, we constructed a simplified cell cycle system and performed SSA (Fig. 2g). Exactly the same results were obtained from the analysis of the simplified system as those from the analysis of the original cell cycle system. This is because the simplified system also contains BS1 and BS2 (Fig. 2g). In addition, the effect of adding Cdc13 monomer degradation is the same as the original network, i.e., BS1 is destroyed, while BS2 is maintained. These behaviors will always appear as long as the following conditions are met: (i) the total amount of chemical A (CDK) is conserved and (ii) chemicals B and C (cyclins) are synthesized, bind to A, and are subsequently dissociated and decomposed. In other words, these behaviors are not specific to the topology of the network we used, and have the potential to universally appear in all possible cell cycle systems. As mentioned above, one of our future goals is to analyze the cell cycle systems of other species using SSA. In many species, the cell cycle is controlled by the dynamics of CDK-cyclin complexes, where the levels of CDKs remain relatively constant, while the levels of cyclins change over time^15^. In such systems, it is highly likely that multiple, non-overlapping buffering structures will appear. We expect that independent control of the cell cycle through buffering structures is universal across species.
3) Third, by iterating between the theoretical predictions and experimental validation, we were even able to update the cell cycle network information that we initially relied on. This was possible because SSA is a topology-based theory that derives predictions from network information alone. If we had used mathematical modeling approaches assuming reaction rate functions and parameter values, a mismatch between theoretical predictions and experimental validation would have required us to consider the possibility that the model assumptions were incorrect. By using SSA, we are able to conclude that discrepancies in the known network information must exist whenever the predicted behavior is not observed experimentally. Furthermore, since SSA can immediately determine the response to any perturbations based on network structure, we are able to comprehensively explore possible network modifications and determine the modifications required to achieve the observed behavior. Our research proposes a new research method for complex network systems in biology. In other words, we can confirm the correctness of the network information and, when necessary, update the information by comparing theoretical predictions based only on network topology with experimental validation results. We believe that we have proposed a new systems biology framework that advances the elucidation of systems through theoretical predictions and experimental validation.

## Acknowledgements

We thank Takashi Okada, Masato Ishikawa, and Yong-Jin Huang for their helpful discussions and comments on the theoretical analysis. We also thank all members of the Aoki Laboratory for their helpful discussions and assistance with experimental work. This research was supported by the CREST program (grant no. JPMJCR1922, JPMJCR24Q4) of the Japan Science and Technology Agency (JST) (http://www.jst.go.jp/EN/index.html), Grant-in-Aid for Scientific Research on Innovative Areas (grant no. 19H03196, 19H05670), Joint Usage/Research Center program of Institute for Life and Medical Sciences Kyoto University. Y.Y. was supported by JSPS KAKENHI (Grant No. 23K14156). H.S. was supported by JSPS KAKENHI grants (nos. 21J01354 and 22K15115) and the Dr. Yoshifumi Jigami Memorial Fund, The Society of Yeast Scientists. Y.G. was supported by a JST, ACT-X grant (no. JPMJAX22B8), and JSPS KAKENHI grants (nos.19K16050 and 22K15110). K.A. was supported by JSPS KAKENHI (Grant No. JP22H02625), Takeda Science Foundation, and Joint Research of the Exploratory Research Center on Life and Living Systems (ExCELLS). (ExCELLS program No. 22EXC348).

## Author contributions

Y.Y. and A.M. conceived the research and wrote the manuscript. Y.Y. conducted both theoretical and experimental work. Theoretical analysis was supervised by A.M., and experimental work was assisted by H.S. and supervised by Y.G. and K.A. All authors discussed the results and commented on the manuscript.

## Competing interests

The authors declare no competing interests.

## Theoretical methods

### 1. Structural sensitivity analysis (SSA) and a buffering structure

#### 1-1: The dynamics of a chemical reaction system

We consider a chemical reaction system^3–5,7^ consisting of *M* chemicals

(*X*_1_, …, *X*_*M*_) and *N* reactions (*R*_1_, ⋯, *R*_*N*_), where reaction *R*_*n*_ is:

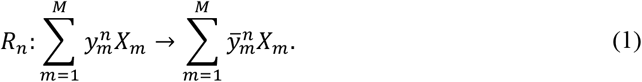

The coefficients 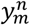 and 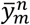 are the numbers of chemical *X*_*m*_ participating in the *n*- th reaction *R*_*n*_ at reactant and product stages, respectively. Dynamical changes in the concentration of chemicals are described by a system of differential equations:

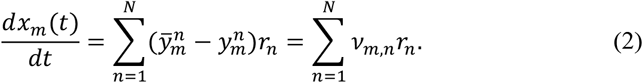

Here *x*_*m*_ (*m* = 1, ⋯, *M*) is the concentration of chemical species *X*_*m*_. The reaction rate (flux) of reaction *R*_*n*_ is *r*_*n*_(*k*_*n*_; ***x***), which depends on the reaction parameter *k*_*n*_ and the chemical concentrations ***x*** ≔ (*x*_1_, ⋯, *x*_*M*_)^*T*^. The stoichiometric coefficient 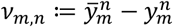 represents the change in the number of the chemical *m* after one reaction *R*_*n*_ occurs.

Equation (2) can be written as:

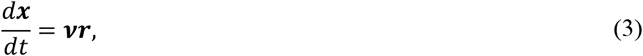

where ***r*** ≔ (*r*_1_, ⋯, *r*_*N*_)^*T*^, and **ν** is the stoichiometric matrix whose *m, n*-th entry is ν_*m,n*_.

The specific form of the reaction rate function *r*_*n*_ is usually unknown, but it is generally accepted that *r*_*n*_ is an increasing function of the concentrations of the substrates of the reaction. Thus, we assume that

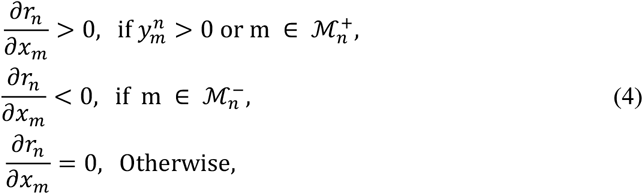

where 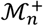 and 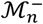 are the set of chemicals that are not a substrate of the reaction *R*_*n*_ but regulate the reaction positively and negatively, respectively. Popular kinetics, such as the mass-action and the Michaelis-Menten kinetics, satisfies this condition.

In addition, each reaction rate function *r*_*n*_ depends on its own reaction parameter *k*_*n*_, which can be the activity or the amount of the enzyme catalyzing reaction *R*_*n*_:

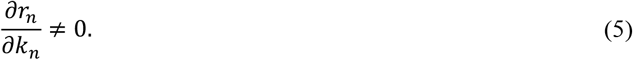

Let ***d***^*l*^ (*l* = 1,…,*L*) be the basis vectors of ker(***v***^*T*^). Since 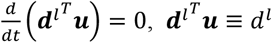 is a conserved quantity in the system (2). In subsection 2, we discuss the case where there is no conserved quantity in the system, i.e., dim (ker(***v***^*T*^)) = 0. In subsection 3, we state the general cases.

#### 1-2: Structural sensitivity analysis (SSA) (In systems without conserved quantities)^3^

Consider a steady state of the system (2). Here, we consider the case where there is no conserved quantity in the dynamical system, i.e., dim(ker(***v***^*T*^)) = 0. In the steady state,

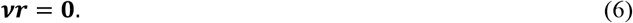

Hence,

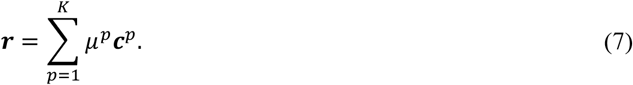

Here, ***c***^*p*^(*p* = 1, ⋯, *K*) is a basis for the null space of ***v***.

We assume that the system reaches a new steady state after a change in enzyme activity 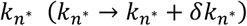.The change in reaction rate of *r*_*n*_ in response to the change is given in the following form using the total derivative:

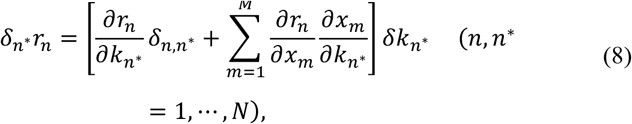

where 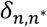 is a Kronecker delta. Throughout this section, all partial derivatives are evaluated at the steady state.

At the same time,

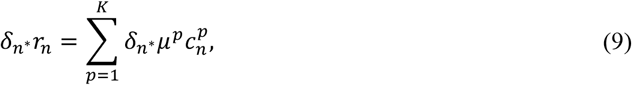

where 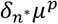 is the change in *μ*^*p*^ due to the change in 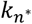.From (8) and (9), we obtain:

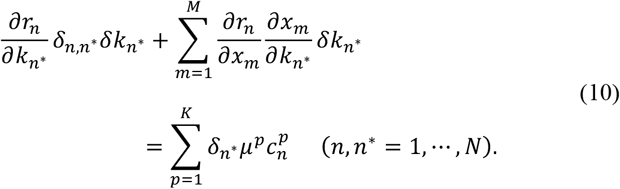

Using 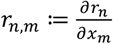 and 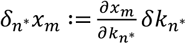,we have:

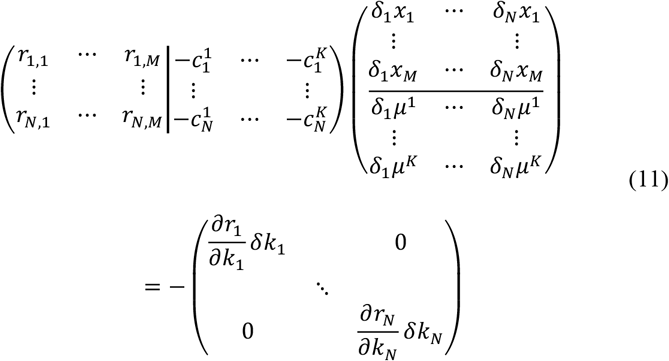

Since dim(ker(***v***^*T*^)) = 0 indicates *M* + *K* = *N*, all matrices that appear in equation (11) are square matrices. Let **A:** = (*r*_*n,m*_|−***c***^*p*^) be the first matrix on the left-hand side of equation (11). The right-hand side is a diagonal matrix, and the diagonal components take nonzero value from equation (5). Assuming that **A** has an inverse matrix, we obtain:

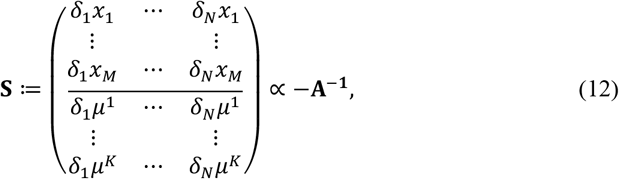

where **S** ∝ −**A**^**−1**^ indicates that **S**_*i,j*_ ≠ 0 ⇔ (−**A**^**−1**^)_*i,j*_ ≠ 0. Each component of **S** shows the response of the steady-state concentration *x*_*m*_ (or that of the steady flow *μ*^*p*^) to the change in the reaction parameter 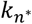.

There are several important things to note about Equation (12).

##### 1. The presence or absence of a response is soley determined by the network structure

The value of *r*_*n,m*_ is nonzero when chemical *m* is a substrate of reaction *R*_*n*_ (or when *m* regulates the reaction rate of *R*_*n*_). Otherwise, *r*_*n,m*_ is constitutively zero. In other words, the pattern of the zero (nonzero) components of the left part of the matrix **A** is determined solely from the structure of the reaction network, and is independent of the choices of the parameters or reaction rate functions. The right part of the matrix **A** (basis vectors for the stoichiometric matrix) is also determined by network structure alone. Hence, zero or nonzero of each component in the matrix **S** ∝ −**A**^**−1**^ is also determined solely from the structure of the reaction network. Since we are interested only in qualitative changes, we let **S** ≔ −**A**^**−1**^ in the following.

##### 2. We can determine the direction of a response, i.e., sign of *δ*_*n*\*_*x*_*m*_ or *δ*_*n*\*_*x*_*n*_

We may be able to determine not only the presence or absence of a response, but also direction of responses, i.e. whether each chemical concentration or reaction rate increases or decreases, under an additional reasonable assumption. For instance, if *k*_*n*_ represents the activity of the enzyme catalyzing the reaction *R*_*n*_, we can assume that 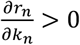,for all *n*. In that case, from (11), the signs of each entry in **S** is same as the signs of responses. Hereafter, unless otherwise specified, we assume that 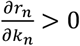,hence the signs of each entry of **S** is the direction of changes upon an increase in each *k*_*n*_.

##### 3. The response pattern can remain valid even for large perturbations

It may seem that equation (12) applies only to small changes in the enzyme activity 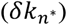.However, a large perturbation can be regarded as a sum of a repeated small perturbations. Since the pattern of zero (non-zero) components in **S** does not depend on on parameter ***k*** or the steate ***x***, it will be always the same, even after repeated changes in 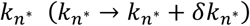.Thus, the response pattern does not change even if a perturbation is large. This holds true as long as there exists a steady state after the perturbation and the steady states are continuous with respect to 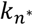 during the repeated changes.

##### Examples

The chemical reaction system in Fig. 1b consists of 3 reactions:

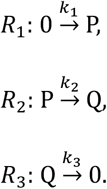

The stoichiometric matrix ***v*** is given by

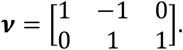

Since ker(***v***) is spanned by (1,1,1)^*T*^ and ker(***v***^***T***^) = {**0**}, the matrix **A** is computed as

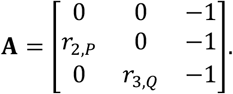

By inverting the matrix **A**,

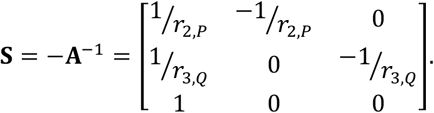

Since *r*_2,*P*_ and *r*_3,*Q*_ are positive, the signs of the entries of **S** are determined as

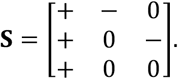

The second column of **S** suggests that an increase in *k*_2_ results in a decrease in the steady-state concentration of chemical P, but the steady-state concentration of chemical Q is not affected (Fig. 1b,c,d). The reaction rate vector at the steady state ***r*** ≔ (*r*_1_, *r*_2_, *r*_3_)^*T*^ is given by ***r*** = *μ*^1^(1,1,1)^*T*^. The last row of the second column indicates that an increase in *k*_2_ does not affect *μ*^1^, and therefore, it does not alter the reaction rates of any reactions in the system (Fig. 1c,d).

#### 1.3: SSA for a system with conserved quantities^5^

So far, we have assumed that the dimension of the left null space of ***v*** is 0, that is, there is no conserved quantity in the system, but we are able to analyze the systems with conserved quantities in the following way^5^. In the presence of conserved quantities, steady-state concentrations are affected not only by reaction parameters but also by the initial values of conserved quantities. Therefore, in this case, there are two types of perturbations: the perturbation of the reaction parameter 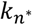 and that of the conserved quantity *d*^*l*^.

We define the **A** matrix as:

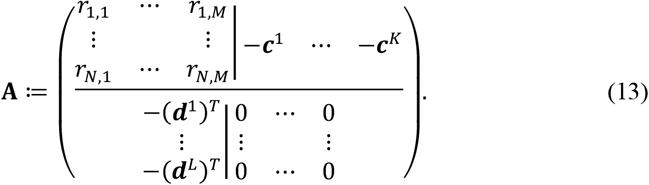

Since *M* + *K* = *N* + *L*, **A** is square. By calculating the inverse of this matrix **A**, the response of the system to the perturbation of parameters is determined at once:

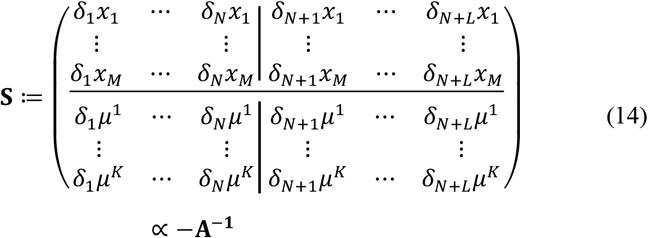

Here, *δ*_*N*+*l*_*x*_*m*_ is the change in the steady state concentration of the chemical *m* when the conserved quantity *d*^*l*^ is perturbed. The *δ*_*N*+*l*_*μ*^*n*^is the change in the steady flow along the basis vector ***c***^*n*^ when *d*^*l*^ is perturbed. Further details of the analysis are as previously described^5^.

##### Examples

The chemical reaction system in Extended Data Fig. 1c consists of 4 reactions

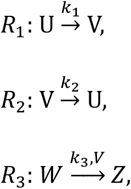

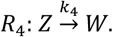

Above the arrow of each reaction *R*_*n*_, we display *k*_*n*_ and 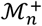.The stoichiometric matrix ***v*** is given by

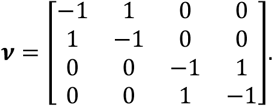

The system has two independent cycles: ker(***v***) =< (1,1,0,0)^*T*^,(0,0,1,1)^*T*^ >. Since ker(***v***^***T***^) =< (1,1,0,0)^*T*^,(0,0,1,1)^*T*^ >, the system has two conserved quantities: *d*^1^ = *x*_*U*_ + *x*_*V*_ and *d*^2^ = *x*_*W*_ + *x*_*Z*_. The matrix **A** is computed as

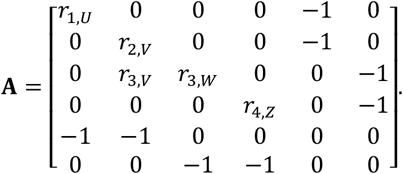

By inverting the matrix **A**, the signs of the entries of **S** are determined.

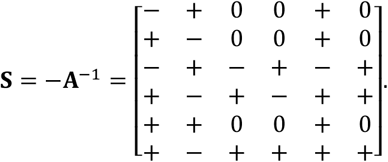

For instance, the third column of **S** indicates that an increase in *k*_3_ results in a decrease in the steady-state concentration of chemical W and an increase in that of Z, but the steady-state concentrations of chemical P and Q are not affected (Extended Data Fig. 1c,d). The last column of **S** indicates that an increase in *d*_2_ results in increases in the steady-state concentrations of chemical W and Z, but the steady-state concentrations of chemical P and Q are not affected.

#### 1-4: SSA for the cell cycle network

The cell cycle system composed of 20 chemicals and 24 reactions^2^. (Fig. 2c). The stoichiometric matrix ***v*** is:

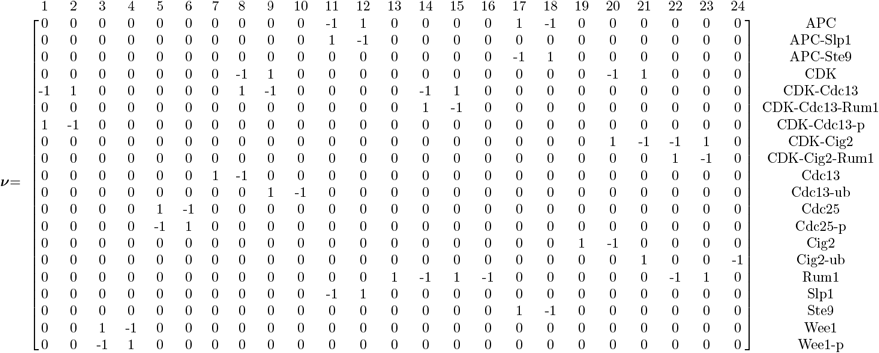

Since *K* = 10, this system has 10 cycles. This system has 6 conserved quantities (*L* = 6):

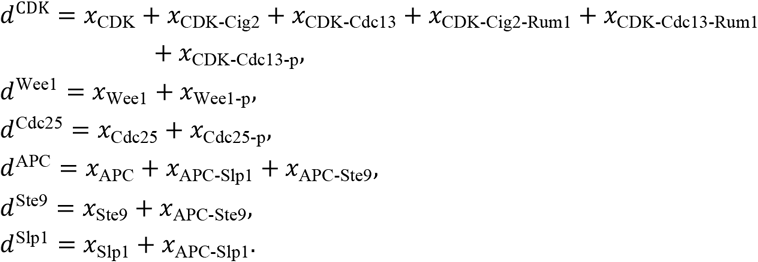

The matrix **A** is computed as:

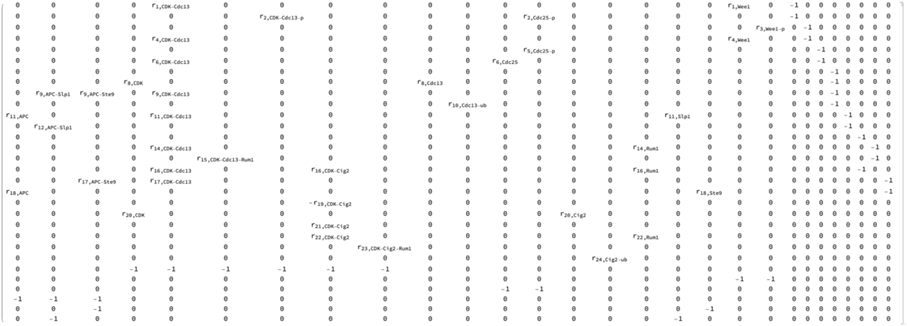

Since regulation of *r* by CDK-Cig2 is negative, we write 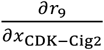 as using the positive variable *r*_19,CDK**−**Cig2_. We calculated **S** = −**A**^**−**1^ wi th the use of Mathematica software. Using *r*_*n,m*_ > 0, the signs of the entries for the upper part of **S** are determined as:

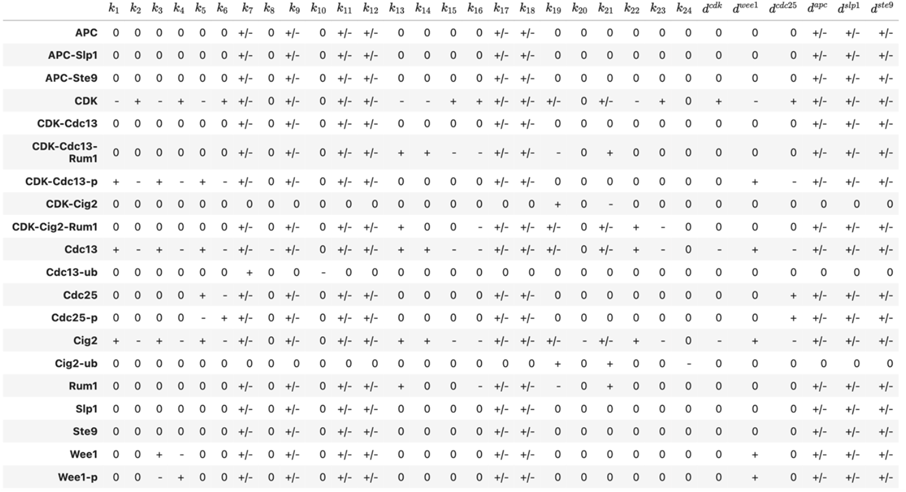

Here, + and − represent qualitative responses to perturbations. The symbol +/− means that the sign depends on quantitative values of *r*_*n,m*_.

Some entries of **S** have undetermined signs (+/−), but assuming the stability of the steady state could allow us to constrain these signs to either + or −. Since asymptotically unstable fixed points cannot exist in biological setting, we can assume that the steady state is asymptotically stable. In a previous study^42^, it was proved that

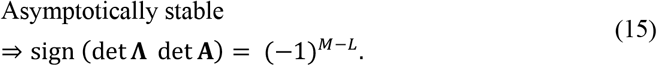

Here,

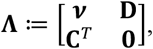

where **C** ≔ [***c***^1^, ⋯, ***c***^*K*^] and **D** ≔ [***d***^1^, ⋯, ***d***^*L*^]. Although the choices of **C** and **D** are not uniquely determined, as long as the same **C** and **D** are used for both **A** and **Λ**, equation (15) holds regardless of the choice of bases. In the cell cycle network, *M* = 20, *L* = 6, and det **Λ** < 0. Therefore, det **A** < 0 is required for the fixed point to be asymptotically stable. The sensitivity **S**_*m,n*_ is given by:

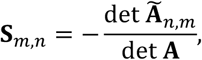

where 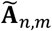 denotes the cofactor of the *m, n*-th element of **A**. If the signs of det 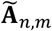 are determined to + or −, the signs of **S**_*m,n*_ is determined accordingly. As a result, we can update the signs of **S** as shown in Table 1. As an example, the steady-state concentration of Cdc2-Cdc13 in response to an increase in *k*_7_ (Cdc13 synthesis rate) was initially undetermined (+/−), but the assumption of asymptotic stability constrains it to +.

#### 1-5: Law of localization and buffering structure

We briefly review the law of localization and buffering structure^4,5^.

##### Definition of a buffering structure

For a given chemical reaction network, we consider a subset of the network Γ = (𝓂, 𝓇), where 𝓂 is a subset of chemical species and 𝓇 is a subset of reactions. We call Γ a buffering structure if Γ satisfies the following two conditions:

1. 𝓇 encompasses all reactions whose reaction rates depend on the concentration of the chemicals within 𝓂, i.e., all reaction arrows that leave from 𝓂 are included in 𝓇 (“output-completeness”).
2. λ(Γ) = 0.

Here the index λ(Γ) is defined as λ(Γ) ≔ −|𝓂| + |𝓇| − *N*(𝓇) + *N*_*c*_(𝓂). *N*(𝓇) is the number of cycles consisting of reactions in 𝓇. *N*_*c*_(𝓂) is the number of independent conserved quantities including at least one element in 𝓂.

To be precise, *N*(𝓇) ≔ dim{***x*** ∈ ker(***v***) | supp ***x*** ⊂ 𝓇}, where supp ***x***≔{*m*|***x***_*m*_ ≠ 0}. We can compute *N*(𝓇) as *N*(𝓇) = dim(ker(***vI***^𝓇^)), where ***I***^𝓇^ ∈ ℝ^*N*×*N*^ is the projection matrix onto 𝓇, i.e., 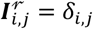 for *i, j* ∈ 𝓇 and 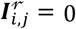 otherwise. We can compute *N*_*c*_(𝓂) as 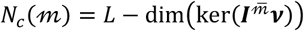,where 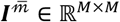 is the projection matrix onto the complement of 𝓂, i.e., 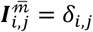 for *i, j* ∉ 𝓂 and 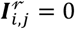 otherwise.

##### Law of Localization^4,5^

The steady-state chemical concentrations and reaction fluxes outside of a buffering structure Γ = (𝓂, 𝓇) do not change under any perturbation of the reaction rate parameters in 𝓇 (or conserved quantities containing at least one chemical in 𝓂).

##### Outline of Proof

Since Γ is an output-complete, we can construct a block diagonal matrix with the elements related to Γ = (𝓂, 𝓇) gathered in the upper left corner by appropriately permutating the rows and columns of **A**:

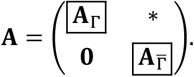

Each row of 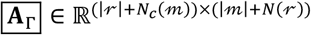 corresponds to a reaction in 𝓇 or a conserved quantity that contains at least one chemical in 𝓂, and each column corresponds to a chemical in 𝓂 or a cycle consisting only of reactions in 𝓇. Since 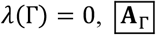,is a square block. Hence,

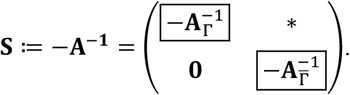

The left lower block of **S** is **0**, which means that any perturbation to a parameter in Γ does not affect any of the chemicals in 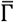.

##### An Example of a buffering structure

In the chemical reaction system in Fig. 1b, Γ = {P, *R*_2_} is a buffering structure (Fig. 1b and Extended Data Fig. 1a,b), since

1. Γ is output-complete (the reaction arrow that leave from P is included in Γ),
2. λ(Γ) = 0 (|𝓂| = 1, |𝓇| = 1, *N*(𝓇) = 0, *N*_*c*_(𝓂) = 0).

##### Finding all buffering structures in a chemical reaction system

In principle, buffering structures can be searched by checking the two conditions (output-completeness and λ(Γ) = 0) for all possible subnetworks. However, the time complexity of this method is *O*(2^*M*+*N*^), which becomes extremely high for large networks. We previously proved that the concept of buffering structure is equivalent to the concept of a regulatory module, as defined below (“the inverse of the law of localization”)^6,47^.

###### Definition of a regulatory module

For a given chemical reaction network, we consider a subset of the network Γ = (𝓂, 𝓇), where 𝓂 is a subset of chemical species and 𝓇 is a subset of reactions. We call Γ a regulatory module if Γ satisfies the following two conditions:

1. Γ is output-complete.
2. Any perturbations in reaction parameters or conserved quantities in Γ does not affect the steady-state concentrations of chemicals outside of 𝓂.

The law of localization states that if subnetwork is a buffering structure, it is a regulatory module. In contrast, the inverse theorem states the opposite: if a subnetwork is a regulatory module, it is a buffering structure. Based on the inverse theorem, finding buffering structures is equivalent to finding regulatory modules, which is achieved through the following procedure^6,47^.

First, we determine whether each component in the matrix **S** ≔ −**A**^**−1**^ is zero or nonzero.

Then, for each reaction *R*_*n*_ (or the conserved quantity *d*^*l*^),

1. Start by constructing a subnetwork that contains only *R*_*n*_ (or *d*^*l*^).
2. Add chemicals whose steady-state concentrations are influenced by the perturbation of *k*_*n*_ (or *d*^*l*^) to the subnetwork.
3. Add the reactions that leaves from the chemicals added in step 2 so that the subnetwork becomes output-complete. (Conserved quantities that contain the chemicals added in step 2 are also added to the subnetwork.)
4. Repeat steps 2 and 3 until no further reactions or chemicals needs to be added.

The resulting subnetwork is a regulatory module and thus is a buffering structure. Details of the algorithm are provided in our previous paper^6^. We conducted this analysis using our previously developed Python package *ibuffpy* (https://github.com/hishidagit/SSApy).

#### 1-6: Hierarchy graph

The nonzero response patterns under perturbations of different parameters can exhibit inclusion relations among them, i.e., exhibit hierarchical structures^3–6^. This hierarchy encompasses every possible perturbation-response pattern. We also found that the hierarchy of perturbation-response patterns corresponds to that of buffering structures. The hierarchy is represented by a “hierarchy graph”, which is a directed acyclic graph *G* with each node *v*_*i*_ consisting of a subset of reaction parameters and chemicals. Key characteristics of *G* are as follows:

1. Each chemical and reaction appears exactly once in *G*.
2. When a reaction rate in a node *v*_*i*_ is perturbed, the chemical in *v*_*i*_, as well as those in *D*_*i*_, show nonzero responses. Here, *D*_*i*_ is the downstream nodes of *v*_*i*_, i.e., *D*_*i*_ ≔{*v*_*j*_}there exists a path from node *v*_*i*_ to *v*_*j*_}. Therefore, reaction parameters that affect a chemical *X* are limited to the reactions in the node containing *X* and those in the upstream nodes.
3. *v*_*i*_ ∪ *D*_*i*_ corresponds to some buffering structure. Furthermore, all buffering structures can be represented as *v*_*i*_ ∪ *D*_*i*_ using some *i* (or a union of them).

Details of the hierarchy are described in our previous paper^6^. The hierarchy graph for the cell cycle network was constructed using our previously developed Python package, *ibuffpy* (https://github.com/hishidagit/SSApy).

### 2. Numerical simulation

In Fig. 1c, reaction rate functions are set as follows: *r*_1_ = *k*_1_, *r*_2_ = *k*_2_*x*_*P*_, *r*_3_ = *k*_3_*x*_*Q*_, with *k*_1_ = *k*_2_ = *k*_3_ = 1.0. The initial conditions are *x*_*P*_ = *x*_*Q*_ = 1.0. At *t* = 12, *k*_2_ is increased by a factor of 2.

In Fig. 2, reaction rate functions are assumed to be combinations of mass action and Michaelis–Menten kinetics:

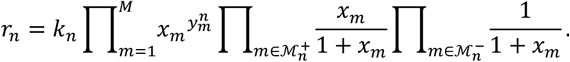

Each reaction parameter *k*_*n*_ is set to 1.0. The initial state is set to be a steady state of the system. At *t* = 150, *k*_19_ (Fig. 2d) or *k*_7_ (Fig. 2e) is increased by a factor of 2.

In Fig. 2f, the numerical simulation of the network dynamics is conducted to recapitulate the cell cycle dynamics. Reaction rate functions, the reaction parameters, and initial values remain the same as those in Fig. 2d,e, except for *k*_19_ and *k*_7_. We assume that the value of *k*_19_ or *k*_7_ change with time, following the equations below:

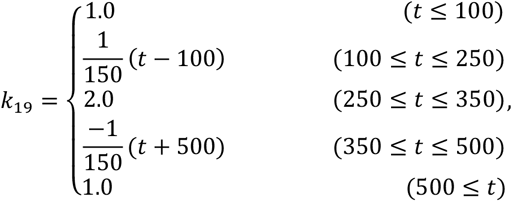

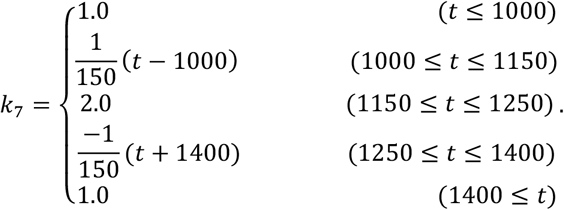

The rates of changes of *k*_19_ or *k*_7_ are slower than the speed at which the system, with fixed parameter values, reaches a steady state. Thus, changes in *k*_19_ or *k*_7_ lead to instantaneous shifts in the steady state concentrations of each chemical. We used the python package scipy.integrate.odient to perform numerical simulation.

### 3. Network modification

Experimental validation demonstrated that BS2 (the buffering structure containing CDK-Cdc13 but not CDK-Cig2) exists, but BS1 (the buffering structure containing CDK-Cig2 but not CDK-Cdc13) does not (Fig. 3). To identify network modifications that account for these experimental results, we considered adding degradation reaction to each chemical that lacks a degradation reaction in the original network (Fig. 4). The chemicals considered for modification included Cdc2, Cig2, Cdc13, CDK-Cdc13, CDK-Cig2, CDK-Cig2-Rum1, CDK-Cdc13-Rum1, APC, Slp1, Wee1, Wee1-p, Cdc25, and Cdc25-p. Then we searched for modifications that break BS1 while maintaining BS2.

Since buffering structures must be output-complete—meaning that all reaction arrows leaving the chemicals in a buffering structure must be included in it—the added degradation reaction of a chemical must be included into buffering structures that contain the chemical. To determine whether a buffering structure is maintained or broken after each modification, we examined the index of BS1 and BS2 defined as λ ≔ −#chemicals + #reactions − #cycles + #conserved quantities. If the index of a buffering structure remains 0, it is maintained; otherwise, it is broken.

| Modifications | BS 1 |  | BS 2 |  |
| --- | --- | --- | --- | --- |
|  | Index | Persistence | Index | Persistence |
| CDK → 0 | $\lambda = 0$ | ✓ | $\lambda = 0$ | ✓ |
| Cig2 → 0 | $\lambda = 0$ | ✓ | $\lambda = 1$ | |
| Cdc13 → 0 | $\lambda = 1$ | | $\lambda = 0$ | ✓ |
| CDK-Cdc13 → 0 | $\lambda = -1$ | | $\lambda = 0$ | |
| CDK-Cig2 → 0 | $\lambda = 0$ | | $\lambda = -1$ | |
| CDK-Cig2-Rum1 → 0 | $\lambda = 0$ | | $\lambda = 0$ | |
| CDK-Cdc13-Rum1 → 0 | $\lambda = 0$ | | $\lambda = 0$ | |
| APC → 0 | $\lambda = 0$ | ✓ | $\lambda = 0$ | ✓ |
| Slp1 → 0 | $\lambda = 0$ | ✓ | $\lambda = 0$ | ✓ |
| Wee1 → 0 | $\lambda = 0$ | ✓ | $\lambda = 0$ | ✓ |
| Wee1-p → 0 | $\lambda = 0$ | ✓ | $\lambda = 0$ | ✓ |
| Cdc25 → 0 | $\lambda = 0$ | ✓ | $\lambda = 0$ | ✓ |
| Cdc25-p → 0 | $\lambda = 0$ | ✓ | $\lambda = 0$ | ✓ |

The gray-highlighted modifications (CDK-Cdc13→0, CDK-Cig2→0, CDK-Cig2-Rum1 →0, CDK-Cdc13-Rum1→0) renders the matrix **A** singular (not invertible). When **A** is singular, structurally stable fixed points do not exist and some chemical concentrations may diverge to infinity, making it unlikely in real biological networks^6^. Thus, we excluded these four modifications from further consideration. Among the remaining modifications, only one modification (Cdc13→0) meets the condition where the index of BS1 changes to a nonzero value while the index of BS2 remains 0, indicating that only this modification breaks BS1 while maintaining BS2.

### Experimental procedures

#### S. pombe strains and cell culture

All fission yeast strains used in this study are summarized in Extended Data Table 1 along with their origins. Unless otherwise noted, the growth medium and other aspects of the experimental procedures for fission yeast were based on those described previously^48^. The transformation protocol was adapted from a previously reported one^49^.

#### Plasmid construction

For fission yeast endogenous tagging, mNeonGreen^50^ and miRFP670^51^ was subcloned into a pFA6a vector using NEBuilder. To generate the fission yeast stable expression plasmid for Cdc13 (or Cdc13-mNeonGreen), Cig2 (or Cig2-mNeonGreen), and Cdc2-miRFP, each gene was subcloned into plasmid for stable knock-in version.2 (pSKI-v2) vectors^52,53^ (Extended Data Table 2).

#### The quantification of the concentrations of the CDK-cyclin complexes using FCCS

To precisely quantify the concentrations of CDK-cyclin complexes under various perturbation conditions, we employed fluorescence correlation spectroscopy (FCS) and fluorescence cross-correlation spectroscopy (FCCS)^27–29,52,54^. FCS and FCCS measure time-series fluorescence intensity within a small confocal volume (∼1 fl). FCS examines the autocorrelation function of temporal fluorescence fluctuation, making it possible to determine the number of fluorescent molecules within the confocal volume. FCCS, on the other hand, utilizes the cross-correlation function between two fluorescent species to infer the extent to which they form a complex, thereby enabling quantification of the complex concentration.

The quantification of the CDK-cyclin complexes concentrations using FCCS was performed as previously described^27^. It was shown that mNeonGreen (mNG) and phycocyanobilin-bound miRFP670 represent a suitable pair for FCCS in living cells from the viewpoint of high photostability and low bleed-through^27^. Thus, we used *cdc2-miRFP; cig2-mNeonGreen* cells to measure the concentration of *CDK-Cig2* and *cdc2-miRFP; cdc13-mNeonGreen* cells to measure the concentration of CDK-Cdc13. To examine the effects of various perturbations on the concentrations of the CDK-cyclin complexes, we used pSKI-v2, which contains a series of promoters and their mutants to drive a wide range of expression levels^53^. We overexpressed cig2-mNeonGreen, cdc13, or cdc2-miRFP in *cdc2-miRFP; cig2-mNeonGreen* cells. Similarly, we overexpressed cdc13-mNeonGreen, cig2, or cdc2-miRFP in *cdc2-miRFP; cdc13-mNeonGreen* cells (Table S2).

FCS data were obtained using a Leica SP8 Falcon confocal microscope (DMI8; Leica) equipped with an objective lens, a HC PL APO 63×/1.20 W motCORR CS2 objective. A white light pulsed laser was used to illuminate the samples, which were used to measure the cell length. Fission yeast cells expressing mNG were measured at 488 nm (laser power 2%) excitation and 490–600 nm emission, and cells expressing miRFP670 were measured at 633 nm (laser power 0.75%) excitation and 640–795 nm emission. Time-series data of fluorescence fluctuations were obtained for 30 seconds.

The structural parameter and effective confocal volume were calibrated using 1 μM Rhodamine 6G (TCI, R0039) in DDW based on the result that the diffusion constant of Rhodamine 6G in DDW is 414 µm^2^/s at room temperature^28,55^. The Rhodamine 6G solution was measured at 520 nm excitation and 550–700 nm emission.

#### Data analysis for FCCS

All data processing and analysis were performed as previously described^27–29,52^. We use Python 3.12.4, with key libraries including NumPy for numerical operations, Pandas for data handling, Matplotlib for plotting, SciPy for signal processing and curve fitting, and the multipletau package for efficient computation of correlation functions.

The recorded fluorescence signals were used to calculate auto- and cross-correlation functions. The fluorescence time series data were first detrended and rescaled for photobleaching correction prior to subsequent analysis. We employed a detrending method based on a combination of moving average and Savitzky-Golay smoothing. For each fluorescence trace, a moving average with a window of 1500 data points was applied to remove high-frequency noise. The sparsely sampled moving-averaged trace was then smoothed using the Savitzky-Golay filter with a window length of 1501 points. This smoothed trace was then used to create an interpolation function that provided a continuous, smoothed baseline across the entire time series. Each original fluorescence trace was corrected by normalizing it by the square root of the ratio of the smoothed baseline to its initial value at time zero. This normalization helps maintain the overall shape of the fluorescence decay while correcting for slow photobleaching trends. The detrended traces were then resmoothed using the same moving average window to prepare them for correlation analysis.

FCS and FCCS analyses were performed on the detrended fluorescence traces.

The auto-correlation function of each channel and the cross-correlation function between the two channels were computed using the multipletau algorithm, which is efficient for calculating correlation functions over logarithmically spaced time delays. The auto-correlation function and the cross-correlation function G(τ) for each channel were modeled using a function that accounts for diffusion dynamics:

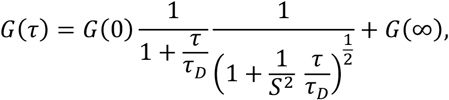

where *τ* is the lag time, *τ*_*D*_ is the diffusion time, 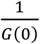 is the average number of fluorescent molecules in the observation volume, and s is the structure parameter. The correlation functions were fitted to these models using nonlinear least-squares optimization to extract the diffusion times and the average number of molecules. The fitting process was constrained by initial estimates and bounded by physically reasonable limits. The concentrations of mNeonGreen (Cig2 or Cdc13), miRFP (CDK), and the complex (CDK-Cig2 or CDK-Cdc13) are given as

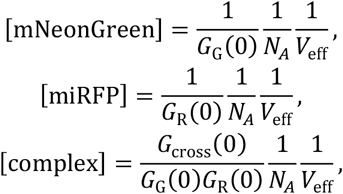

where *G*_G_(τ), *G*_R_(τ), and *G*_cross_(τ) denote the autocorrelation functions of mEGFP, miRFP, and the cross-correlation function, *N*_*A*_ corresponds to the Avogadro number and *V*_eff_ is the effective confocal volume, determined as described above.

When measuring CDK-Cdc13, we selected cells longer than 10.5 μm to identify cells near the G_2_-M transition. We did this selection because, in the early stages of the cell cycle, when Cdc13 expression is low, it is difficult to detect a reliable mNG fluorescence signal or obtain accurate FCCS data. Additionally, CDK might be highly phosphorylated before the G2-M transition, meaning that FCCS results might not accurately reflect the concentration of CDK-Cdc13, as they could include the concentration of CDK-Cdc13-p. Similarly, when measuring CDK-Cig2, we selected cells shorter than 10.5 μm to identify cells near the G_1_-S transition.

#### Time-lapse cycloheximide (CHX) chase

CHX chase analysis of Cdc13 in the fission yeast cells was conducted with reference to previous studies^19,32^. While previous studies performed CHX chase analysis on bulk cell populations, we employed time-lapse live-cell imaging to observe the fluorescence signals of *cdc2-miRFP; cdc13-mNeonGreen* cells following the CHX addition. This approach enables us to measure the degradation rate of Cdc13 separately for cells before and after nuclear division. For a negative control, we used cells expressing mNeonGreen under the control of the *adh1* promotor. The ONIX Microfluidic platform (Merck) was used for time-lapse live-cell imaging of fission yeast cells as previously described^53^. Briefly, growing fission yeast cells were concentrated by centrifugation (3000 rpm, 1 min) and resuspended in 100 mL of the appropriate medium. The suspension was loaded into a trapping chamber (Y04T-04) by imposing pressure of 8 psi for 15 sec. After 3 hours of the culture without drugs, 100 ug/ml cycloheximide, with or without 50 μM Apcin (MedChemExpress), is added. During the imaging, the media was perfused by imposing pressure of 1 psi.

#### Image analysis and data visualization

All the images in CHX chase analysis (Fig. 5) were analyzed using Fiji/ImageJ (https://fiji.sc/) as previously described^27,53^. The background was subtracted by the rolling-ball method (rolling ball radius, 50.0 pixels) adopted in Fiji. An optimized tracking plugin for Fiji, LIM Tracker^56^ enabled easy single-cell tracking. Regions of interest (ROIs) typically corresponded to the nucleus, as determined from miRFP images. Each fluorescence intensity was defined as the mean value of ROIs. We distinguish cells that undergo nuclear division during the tracking (cells in M phase) and cells that do not (cells in G_2_ phase). Accumulated numerical data were visualized using Python with seaborn and matplotlib.

#### Plot and statistical analysis

The box-and-whisker plots in the manuscript show the quartiles of the accumulated data with whiskers denoting the minimum and maximum except for the outliers detected as 1.5 times the interquartile range. The statistical tests used in each plot are listed below: Two-sided Student’s t-test (Fig. 4b, c, e, g, h, j); Student’s t test adjusted by Bonferroni method (Fig. 5e).

---

**Extended Data Fig. 1.**
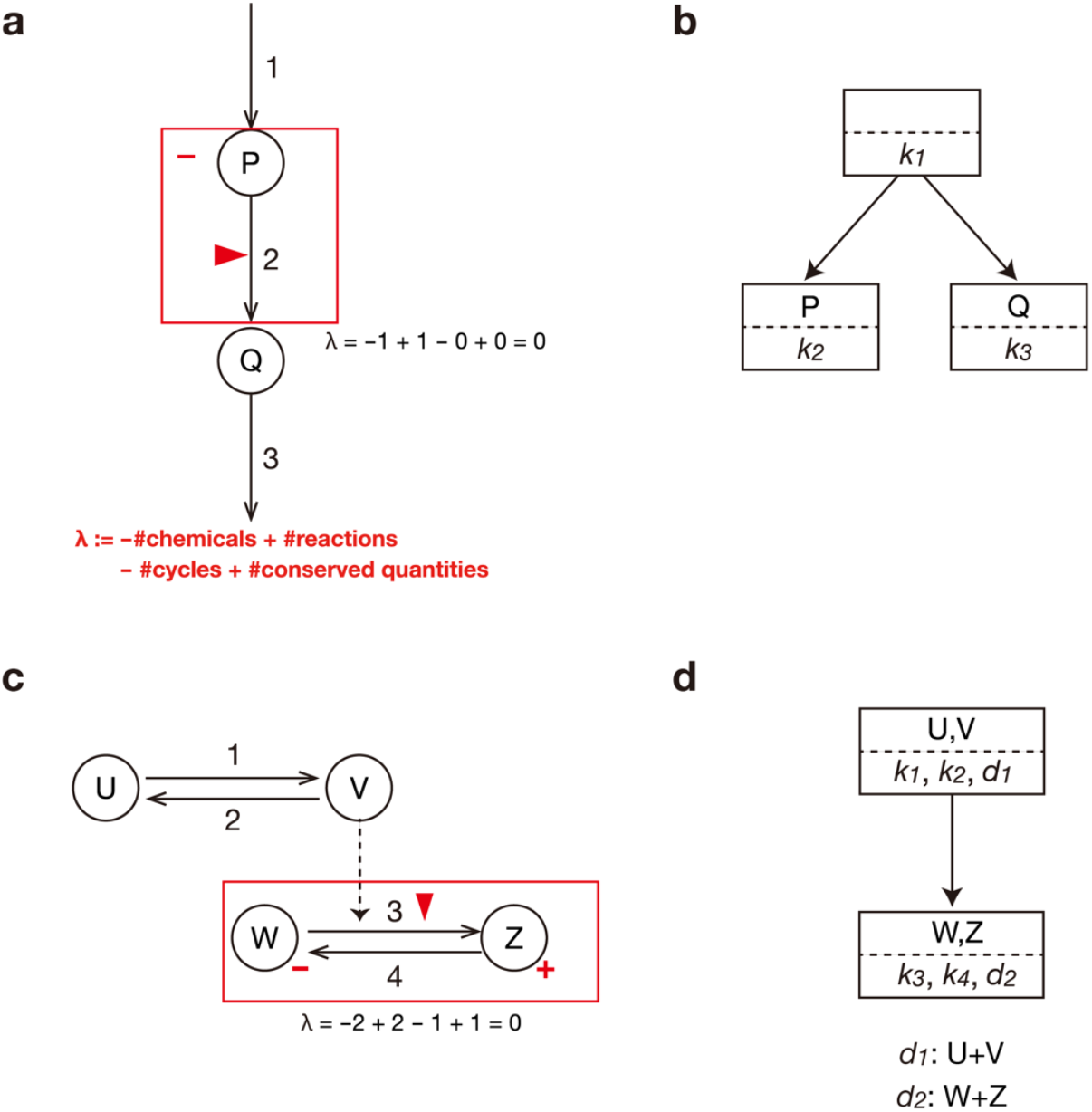
Buffering structure determines the extent to which changes in the parameters propagate within the network. **a**, Graphical representation of a chemical reaction system, as in Fig. 1b, shown here again. Solid lines indicate chemical reactions. The red triangle indicates the reaction that is activated. The signs (increase/decrease) of responses are represented by +/− for chemicals, while chemicals without signs remain unchanged. The subnetwork enclosed by a red box is a buffering structure, with the index λ being equal to 0. **b**, A graph of response hierarchy, which summarizes the inclusion relations between nonzero response patterns in **a**. Modulating the enzyme activity of reactions within a square box leads to nonzero responses in the chemicals within that box and those in the lower boxes, leaving the other chemicals unaffected. **c**, Graphical representation of a chemical reaction system comprising four chemicals (U,V,W,Z) and four reactions (1,2,3,4) with two conserved quantities (*d*_1_:U+V and *d*_2_:W+Z). Solid lines indicate chemical reactions, while the dashed line indicates active regulation. Notations align with those in **a**. The subnetwork enclosed by a red box is a buffering structure, with the index λ being equal to 0. **d**, A graph of response hierarchy in **c**.

**Extended Data Fig. 2.**
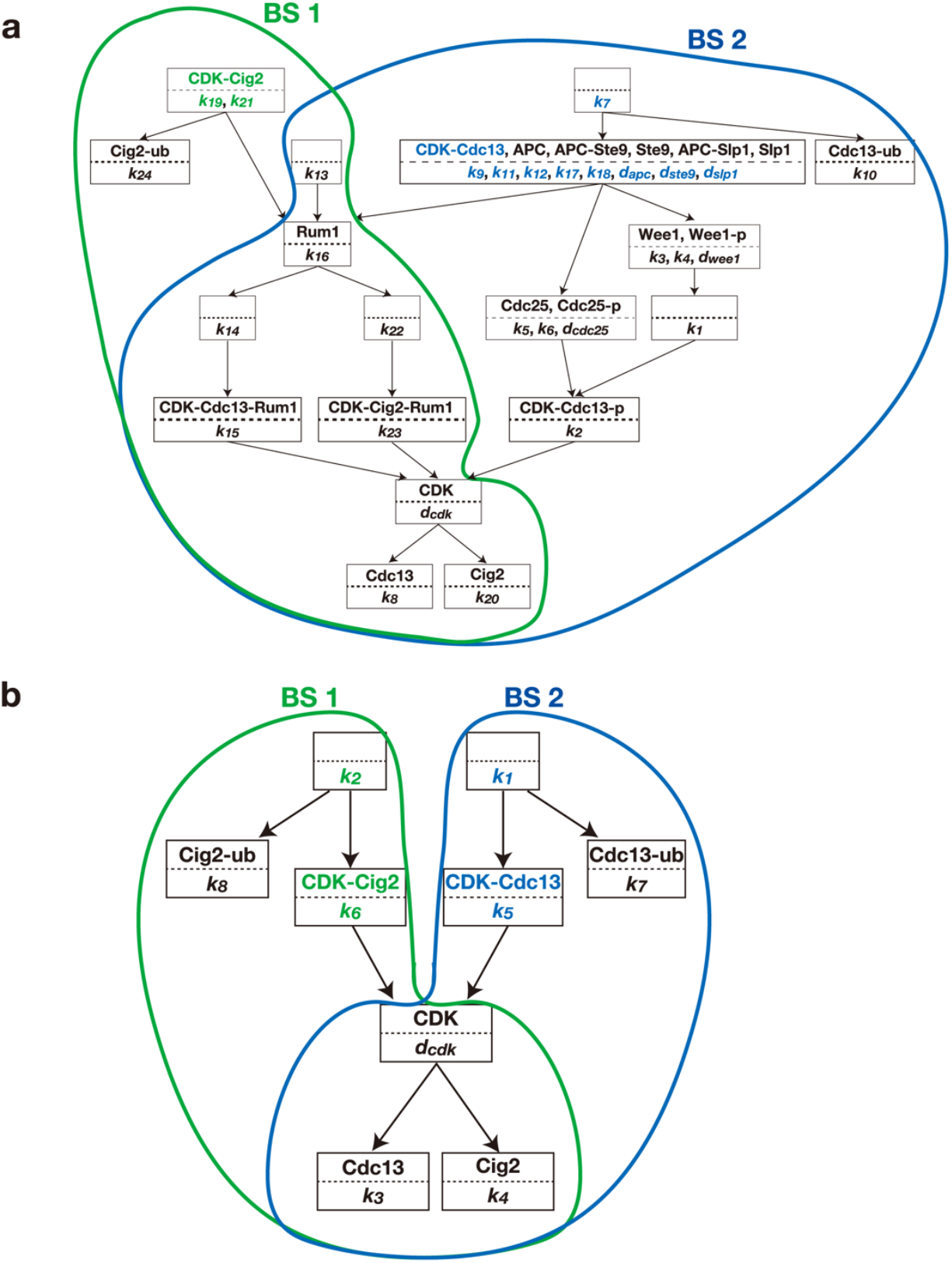
Hierarchy of buffering structures in the cell cycle network. **a**, A graph of response hierarchy of the cell cycle network (Fig. 2c), which summarizes the inclusion relations between nonzero response patterns. Modulating the enzyme activity of reactions within a square box leads to nonzero responses in the chemicals within that box and those in the lower boxes, leaving the other chemicals unaffected. The reaction rate parameters that affect CDK-Cig2 (green) are those located within the box containing CDK-Cig2 or its upstream boxes. The reaction rate parameters that affect CDK-Cdc13 (blue) are those located within the box containing CDK-Cdc13 or its upstream boxes. Subnetworks enclosed by green and blue represent BS1 and BS2, respectively. **b**, A graph of response hierarchy of the simplified cell cycle network (Fig. 2g).

**Extended Data Fig. 3.**
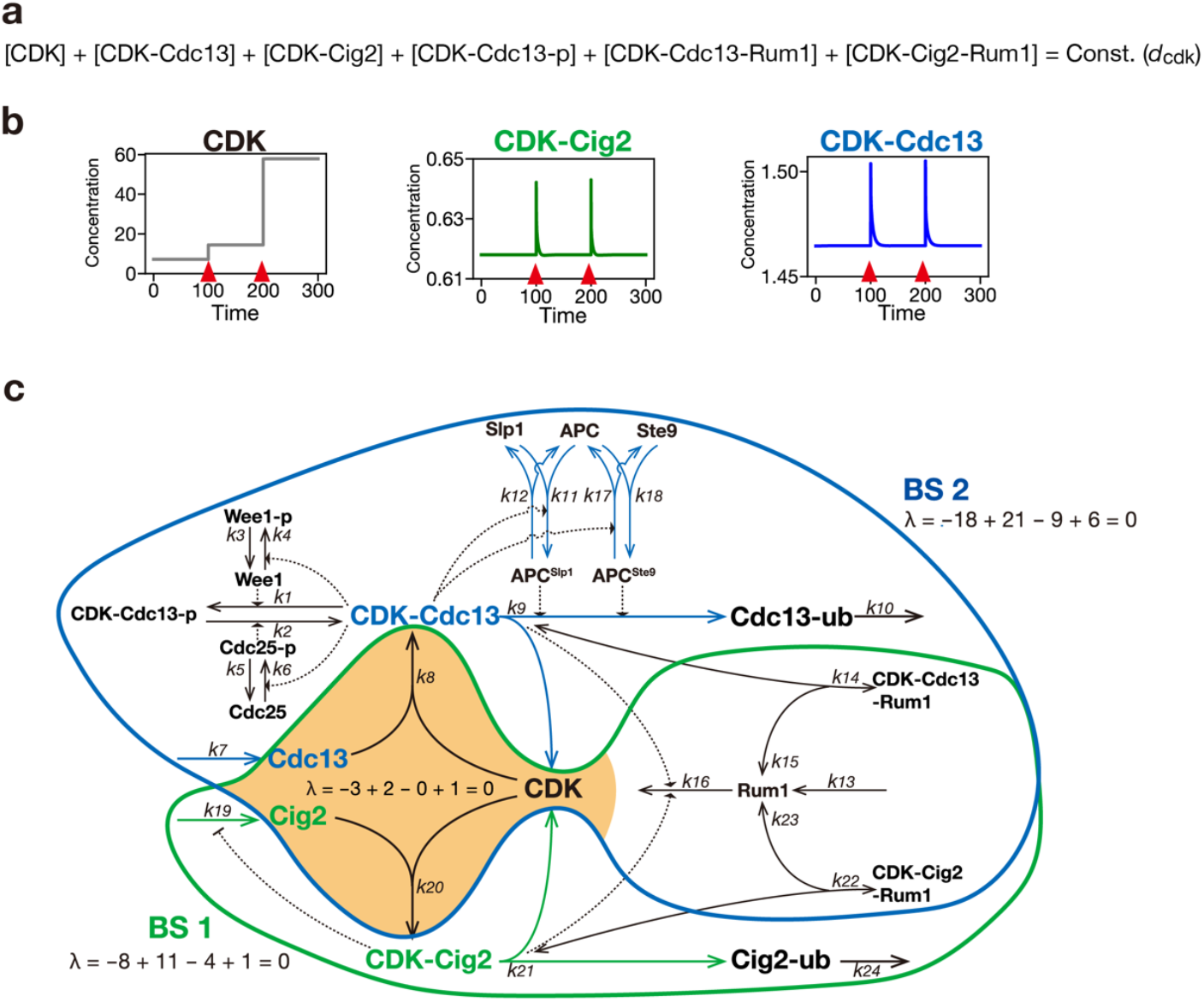
The total CDK amount is predicted to influence neither CDK-Cig2 nor CDK-Cdc13. **a**, The total amount of CDK is constant throughout the dynamics. **b**, The subnetwork highlighted in orange is a buffering structure, which is included by the intersection between BS1 (green) and BS2 (blue). This buffering structure contains CDK, a component of a conserved quantity *d*_cdk_. Thus, the perturbation to the total CDK amount does not affect either CDK-Cig2 nor CDK-Cdc13, both of which is located outside of the buffering structure. **c**, Numerical calculation of the network dynamics are performed. Reaction rate functions are set in the same way as in Fig. 2d,e. At *t* = 100 and *t* = 200, the amount of free CDK, which does not form complexes with other proteins, is doubled.

**Extended Data Fig. 4.**
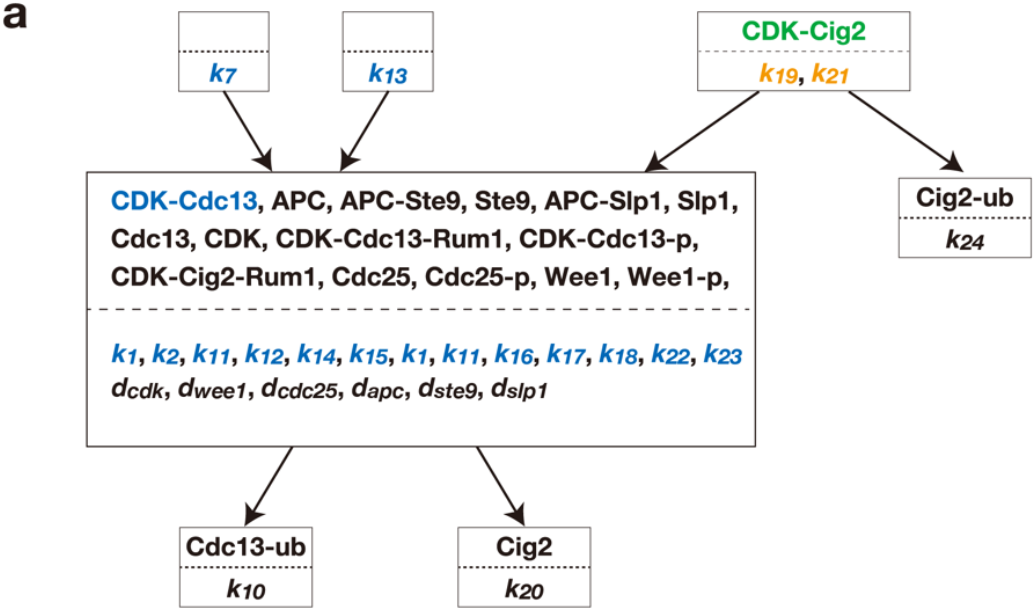
Theoretically identifying missing reactions in the cell cycle network. **a**, The hierarchy graph of the corrected network in Fig. 4b, where Cdc13 that does not bind to CDK is degraded. Reaction rate parameters colored in blue affect CDK-Cdc13 but not CDK-Cig2. Reaction rate parameters colored in orange affect both CDK-Cig2 and CDK-Cdc13.

**Extended Data Fig. 5.**
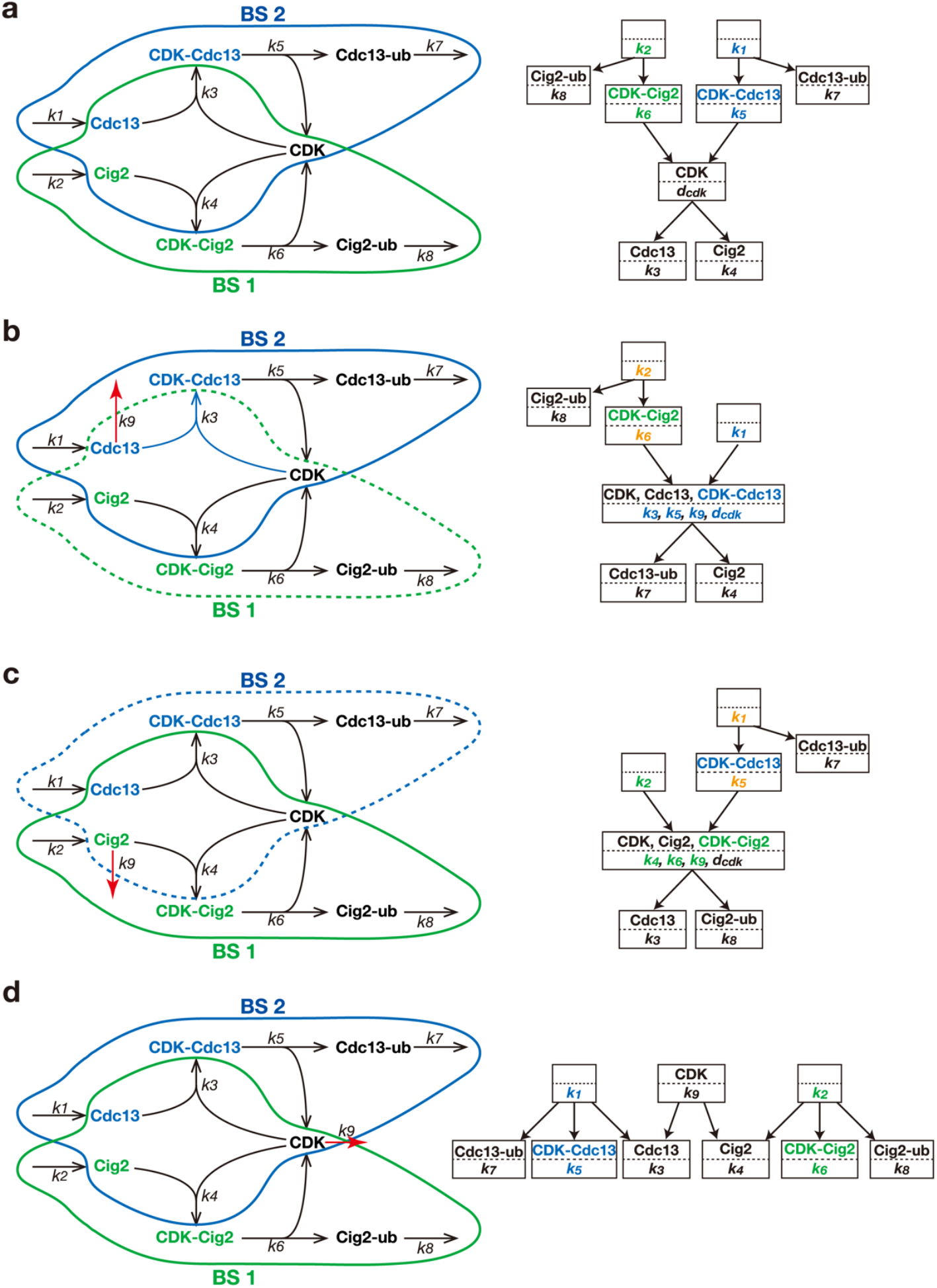
Theoretically identifying missing reactions in the simplified cell cycle network. **a**, The simplified cell cycle network (same as Fig. 2g). The green and blue curve represents BS1 and BS2, respectively. The hierarchy graph is shown in the right. **b-d**, Degradation reactions are introduced for each chemical that lacks a degradation reaction in the original network **a**. Only **b** breaks BS1 while preserving BS2. Reaction rate parameters colored in green affect CDK-Cig2 but not CDK-Cdc13. Reaction rate parameters colored in blue affect CDK-Cdc13 but not CDK-Cig2. Reaction rate parameters colored in orange affect both CDK-Cig2 and CDK-Cdc13.

**Extended Data Table 1.** Fission yeast strain list.

| Strain name | Genotype | Figure | Source |
| --- | --- | --- | --- |
| HS494' (LabID) | h- 1L::Padh1-SynPCB2.1<<bsd cdc2-miRFP670<<hyg cdc13-spmNeonGreen<<kan | Fig. 3,5 | Toyama et al., |
| HS618 (LabID) | h- 1L::Padh1-SynPCB2.1<<bsd cdc2-miRFP670<<hyg cdc13-spmNeonGreen<<kan 2L::Ptdh15-cdc13-spmNeonGreen<<nat | Fig. 3 | This study |
| HS620 (LabID) | h- 1L::Padh1-SynPCB2.1<<bsd cdc2-miRFP670<<hyg cdc13-spmNeonGreen<<kan 2L::Ptdh15-cig2<<nat | Fig. 3 | This study |
| HS622 (LabID) | h- 1L::Padh1-SynPCB2.1<<bsd cdc2-miRFP670<<hyg cdc13-spmNeonGreen<<kan 2L::Padh11-cdc2-miRFP670<<nat | Fig. 3 | This study |
| HS495 (LabID) | h- 1L::Padh1-SynPCB2.1<<bsd cdc2-miRFP670<<hyg cig2-spmNeonGreen<<kan | Fig. 3 | This study |
| HS674 (LabID) | h- 1L::Padh1-SynPCB2.1<<bsd cdc2-miRFP670<<hyg cig2-spmNeonGreen<<kan Padh13-cig2-mNeonGreen<<nat | Fig. 3 | This study |
| HS616 (LabID) | h- 1L::Padh1-SynPCB2.1<<bsd cdc2-miRFP670<<hyg cig2-spmNeonGreen<<kan 2L::Ptdh15-cdc13<<nat | Fig. 3 | This study |
| HS608 (LabID) | h- 1L::Padh1-SynPCB2.1<<bsd cdc2-miRFP670<<hyg cig2-spmNeonGreen<<kan 2L::Padh11-cdc2-miRFP670<<nat | Fig. 3 | This study |
| HS565 (LabID) | h- 2L::Padh1-spmNeonGreen<<nat 3R::Padh1-miRFP670<<kan 1L::synPCB2.1<<bsd | Fig. 5 | Toyama et al., |

**Extended Data Table 2.**
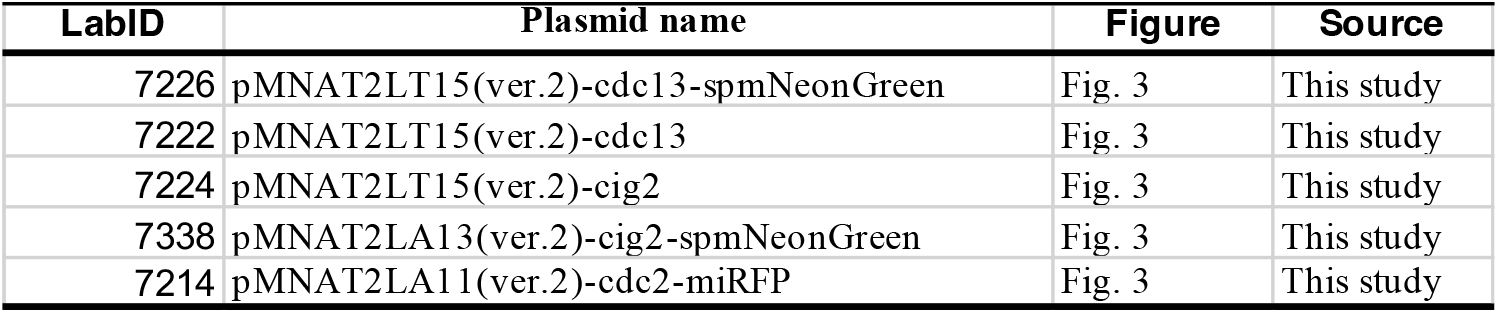
Plasmid list.

| LabID | Plasmid name | Figure | Source |
| --- | --- | --- | --- |
| 7226 | pMNAT2LT15(ver.2)-cdc13-spmNeonGreen | Fig. 3 | This study |
| 7222 | pMNAT2LT15(ver.2)-cdc13 | Fig. 3 | This study |
| 7224 | pMNAT2LT15(ver.2)-cig2 | Fig. 3 | This study |
| 7338 | pMNAT2LA13(ver.2)-cig2-spmNeonGreen | Fig. 3 | This study |
| 7214 | pMNAT2LA11(ver.2)-cdc2-miRFP | Fig. 3 | This study |

## Notes

### Competing Interest Statement

The authors have declared no competing interest.

## References

1. Sveiczer, A., Csikasz-Nagy, A., Gyorffy, B., Tyson, J. J. & Novak, B. Modeling the fission yeast cell cycle: Quantized cycle times in wee1-cdc25Δ mutant cells. Proc Natl Acad Sci U S A 97, 7865–7870 (2000).

2. Sveiczer, A., Tyson, J. J. & Novak, B. Modelling the Fission Yeast Cell Cycle. https://academic.oup.com/bfg/article/2/4/298/224898.

3. Mochizuki, A. & Fiedler, B. Sensitivity of chemical reaction networks: A structural approach. 1. Examples and the carbon metabolic network. J Theor Biol 367, 189–202 (2015).

4. Okada, T. & Mochizuki, A. Law of Localization in Chemical Reaction Networks. Phys Rev Lett 117, (2016).

5. Okada, T. & Mochizuki, A. Sensitivity and network topology in chemical reaction systems. Phys Rev E 96, (2017).

6. Yamauchi, Y., Hishida, A., Okada, T. & Mochizuki, A. Finding regulatory modules of chemical reaction systems. Phys Rev Res 6, (2024).

7. Feinberg, M. REVIEW ARTICLE NUMBER 25 CHEMICAL REACTION NETWORK STRUCTURE AND THE STABILITY OF COMPLEX ISOTHERMAL REACTORS-I. THE DEFICIENCY ZERO AND DEFICIENCY ONE THEOREMS. Chemical Engineering Science vol. 42.

8. Horn, F. Necessary and Sufficient Conditions for Complex Balancing in Chemical Kinetics.

9. MartinnFeinberg Foundations of Chemical Reaction Network Theory. http://www.springer.com/series/34.

10. Orth, J. D., Thiele, I. & Palsson, B. O. What is flux balance analysis? Nature Biotechnology vol. 28 245–248 Preprint at 10.1038/nbt.1614 (2010).

11. Kodama, M. et al. A shift in glutamine nitrogen metabolism contributes to the malignant progression of cancer. Nat Commun 11, (2020).

12. Ohno, S. et al. Kinetic Trans-omic Analysis Reveals Key Regulatory Mechanisms for Insulin-Regulated Glucose Metabolism in Adipocytes. iScience 23, (2020).

13. Zhao, S. et al. Regulation of cellular metabolism by protein lysine acetylation. Science (1979) 327, 1000–1004 (2010).

14. Ferjani, A. et al. Pyrophosphate inhibits gluconeogenesis by restricting UDP-glucose formation in vivo. Sci Rep 8, (2018).

15. Morgan, D. O. Principles of CDK regulation. Nature 1995 374:6518 374, 131– 134 (1995).

16. Nurse, P., Thuriaux, P. & Nasmyth, K. Genetic Control of the Cell Division Cycle in the Fission Yeast Schizosaccharomyces Pombe. vol. 146 (1976).

17. Stern, B. & Genetics, P. N. A quantitative model for the cdc2 control of S phase and mitosis in fission yeast. Trends in Genetics 12, 345–350 (1996).

18. Ayté, J., Schweitzer, C., Zarzov, P., Nurse, P. & DeCaprio, J. A. Feedback Regulation of the MBF Transcription Factor by Cyclin Cig2. NATURE CELL BIOLOGY vol. 3 http://cellbio.nature.com (2001).

19. Bashir, S. et al. Size-Dependent Expression of the Fission Yeast Cdc13 Cyclin is Conferred by Translational Regulation. doi:10.1101/2023.01.16.524304.

20. Curran, S., Dey, G., Rees, P. & Nurse, P. A quantitative and spatial analysis of cell cycle regulators during the fission yeast cycle. (2022) doi:10.1073/pnas.

21. Moriya, H., Chino, A., Kapuy, O., Csikász-Nagy, A. & Novák, B. Overexpression limits of fission yeast cell-cycle regulators in vivo and in silico. Mol Syst Biol 7, (2011).

22. Novak, B., Pataki, Z., Ciliberto, A. & Tyson, J. J. Mathematical model of the cell division cycle of fission yeast. Chaos vol. 11 277–286 Preprint at 10.1063/1.1345725 (2001).

23. Moreno, S. and N. P. Regulation of progression through the Gl phase of the cell cycle by the rum1+ gene. Nature 367, 236–242 (1994).

24. Gould, K. L. & Nurse, P. Tyrosine Phosphorylation of the Fission Yeast Cdc2+ Protein Kinase Regulates Entry into Mitosis. (1989).

25. Perry, J. A. & Kornbluth, S. Cdc25 and Wee1: Analogous opposites? Cell Division vol. 2 Preprint at 10.1186/1747-1028-2-12 (2007).

26. Hirono, Y., Okada, T., Miyazaki, H. & Hidaka, Y. Structural reduction of chemical reaction networks based on topology. Phys Rev Res 3, (2021).

27. Toyama, A. et al. Quantification of Cyclin-CDK dissociation constants in living cells using fluorescence cross-correlation spectroscopy with green and near-infrared fluorescent proteins. Preprint at 10.1101/2024.11.14.623553 (2024).

28. Sadaie, W., Harada, Y., Matsuda, M. & Aoki, K. Quantitative In Vivo Fluorescence Cross-Correlation Analyses Highlight the Importance of Competitive Effects in the Regulation of Protein-Protein Interactions. Mol Cell Biol 34, 3272–3290 (2014).

29. Komatsubara, A. T., Goto, Y., Kondo, Y., Matsuda, M. & Aoki, K. Single-cell quantification of the concentrations and dissociation constants of endogenous proteins. Journal of Biological Chemistry 294, 6062–6072 (2019).

30. Primorac, I. & Musacchio, A. Panta rhei: The APC/C at steady state. Journal of Cell Biology vol. 201 177–189 Preprint at 10.1083/jcb.201301130 (2013).

31. Hishida, A., Okada, T. & Mochizuki, A. Patterns of change in regulatory modules of chemical reaction systems induced by network modification. PNAS Nexus 3, (2024).

32. Esposito, E. et al. Mitotic checkpoint gene expression is tuned by codon usage bias. EMBO J 41, (2022).

33. Berthet, C., Aleem, E., Coppola, V., Tessarollo, L. & Kaldis, P. Cdk2 Knockout Mice Are Viable. Current Biology 13, 1775–1785 (2003).

34. Malumbres, M. et al. Mammalian Cells Cycle without the D-Type Cyclin-Dependent Kinases Cdk4 and Cdk6 Ing to Histone Deacetylases and Chromatin Remodeling Proteins (Reviewed in Harbour and Dean [2000]; Stevaux and Dyson [2002]). The Complexity of Rb Function Is. Cell vol. 118 (2004).

35. Santamaría, D. et al. Cdk1 is sufficient to drive the mammalian cell cycle. Nature 448, 811–815 (2007).

36. Knudsen, E. S. et al. CDK/cyclin dependencies define extreme cancer cell-cycle heterogeneity and collateral vulnerabilities. Cell Rep 38, (2022).

37. Arora, M. et al. Rapid adaptation to CDK2 inhibition exposes intrinsic cell-cycle plasticity. Cell 186, 2628-2643.e21 (2023).

38. Alon, U. Network motifs: Theory and experimental approaches. Nature Reviews Genetics vol. 8 450–461 Preprint at 10.1038/nrg2102 (2007).

39. Alon, Uri. An Introduction to Systems Biology : Design Principles of Biological Circuits. (CRC Press, 2020).

40. Okada, T., Tsai, J. C. & Mochizuki, A. Structural bifurcation analysis in chemical reaction networks. Phys Rev E 98, (2018).

41. Okada, T., Mochizuki, A., Furuta, M. & Tsai, J. C. Flux-augmented bifurcation analysis in chemical reaction network systems. Phys Rev E 103, (2021).

42. Huang, Y.-J., Okada, T. & Mochizuki, A. Uncovering Bifurcation Behaviors of Biochemical Reaction Systems from Network Topology. Preprint at 10.1101/2024.11.19.624026 (2024).

43. Fiedler, B., Mochizuki, A., Kurosawa, G. & Saito, D. Dynamics and Control at Feedback Vertex Sets. I: Informative and Determining Nodes in Regulatory Networks. J Dyn Differ Equ 25, 563–604 (2013).

44. Mochizuki, A., Fiedler, B., Kurosawa, G. & Saito, D. Dynamics and control at feedback vertex sets. II: A faithful monitor to determine the diversity of molecular activities in regulatory networks. J Theor Biol 335, 130–146 (2013).

45. Kobayashi, K., Maeda, K., Tokuoka, M., Mochizuki, A. & Satou, Y. Controlling Cell Fate Specification System by Key Genes Determined from Network Structure. iScience 4, 281–293 (2018).

46. Kobayashi, K., Maeda, K., Tokuoka, M., Mochizuki, A. & Satou, Y. Using linkage logic theory to control dynamics of a gene regulatory network of a chordate embryo. Sci Rep 11, (2021).

47. Hirono, Y., Gupta, A. & Khammash, M. Complete characterization of robust perfect adaptation in biochemical reaction networks. (2023).

48. Mor∼no, S., Klar, A. & Nurse, P. MOLECULAR BIOLOGY OF THE FISSION YEAST S. Pombe 795 [56] Molecular Genetic Analysis of Fission Yeast. vol. 56.

49. Suga, M. & Hatakeyama, T. A rapid and simple procedure for high-efficiency lithium acetate transformation of cryopreserved Schizosaccharomyces pombe cells. Yeast 22, 799–804 (2005).

50. Steiert, F., Petrov, E. P., Schultz, P., Schwille, P. & Weidemann, T. Photophysical Behavior of mNeonGreen, an Evolutionarily Distant Green Fluorescent Protein. Biophys J 114, 2419–2431 (2018).

51. Shcherbakova, D. M. et al. Bright monomeric near-infrared fluorescent proteins as tags and biosensors for multiscale imaging. Nat Commun 7, (2016).

52. Sakai, K. et al. Near-infrared imaging in fission yeast using a genetically encoded phycocyanobilin biosynthesis system. J Cell Sci 134, (2021).

53. Sugiyama, H., Goto, Y., Kondo, Y., Coudreuse, D. & Aoki, K. Live-cell imaging defines a threshold in CDK activity at the G2/M transition. Dev Cell 59, 545-557.e4 (2024).

54. Bacia, K., Kim, S. A. & Schwille, P. Fluorescence cross-correlation spectroscopy in living cells. Nat Methods 3, 83–89 (2006).

55. Müller, C. B. et al. Precise measurement of diffusion by multi-color dual-focus fluorescence correlation spectroscopy. EPL 83, (2008).

56. Aragaki, H., Ogoh, K., Kondo, Y. & Aoki, K. LIM Tracker: a software package for cell tracking and analysis with advanced interactivity. Sci Rep 12, (2022).

